# Polarization Vision—Overcoming Challenges of Working with a Property of Light We Barely See

**DOI:** 10.1101/207217

**Authors:** James J. Foster, Shelby E. Temple, Martin J. How, Ilse M. Daly, Camilla R. Sharkey, David Wilby, Nicholas W. Roberts

## Abstract

In recent years, the study of polarization vision in animals has seen numerous breakthroughs, not just in terms of what is known about the function of this sensory ability, but also in the experimental methods by which polarization can be controlled, presented and measured. Once thought to be limited to only a few animal species, polarization sensitivity is now known to be widespread across many taxonomic groups, and advances in experimental techniques are, in part, responsible for these discoveries. Nevertheless, its study remains challenging, perhaps because of our own poor sensitivity to the polarization of light, but equally as a result of the slow spread of new practices and methodological innovations within the field. In this review, we introduce the most important steps in designing and calibrating polarized stimuli, within the broader context of areas of current research and the applications of new techniques to key questions. Our aim is to provide a constructive guide to help researchers, particularly those with no background in the physics of polarization, to design robust experiments that are free from confounding factors.

## 1. Introduction

### 1.1 The Challenge of Studying Polarization Sensitivity

Polarized light is abundant in nature and, since the discovery of polarization-sensitive orientation in honeybees (von Frisch, 1949), a great number of animal species have been shown to use polarized light to inform their behaviour (Horváth & Varjú, 2004). While most research, until very recently, has focused on navigation and orientation behaviours that use wide-field environmental polarization cues, we are now beginning to uncover the remarkable complexity of the polarization patterns within visual scenes that many species are able to see and use.

That it has taken us so long to appreciate the value of these polarization cues and signals, and the sensitivity of animals to them, is no doubt due to the limited polarization sensitivity of our own visual system. Although, under the correct circumstances, humans can detect and identify polarized light via the Haidinger’s brushes phenomenon (Haidinger, 1844; Shurcliff, 1955; Temple *et al.*, 2015), we are ‘polarization blind’ in our daily lives when compared with the majority of animal species. It has only been with the aid of recently developed polarization imaging technologies (e.g. Powell & Gruev, 2013; Roberts *et al.*, 2014; York *et al.*, 2014; Gagnon & Marshall, 2016) that we are beginning to uncover the complexities of the polarization patterns that exist in nature and, with this, starting to understand how animals use this information.

The aim of this review is to provide a constructive guide for researchers who wish to study polarization sensitivity, but may be unfamiliar with the terminology (Section 2), the measurement techniques (Section 3), the advantages and drawbacks of different means of producing polarized stimuli (Section 4), and, most importantly, the various pitfalls of intensity and spectral confounds (Section 5). For researchers who are more familiar with the field, this review provides an up-to-date collection of the various methods developed to address these challenges. More broadly, this review may be used as a guidebook in the early stages of planning a study; to help determine the equipment, measurements, and control experiments necessary. Finally, we hope that the discussions below will inspire new research, lead to further improvements to these methods, and aid in the interpretation of animal polarization vision experiments.

### 1.2 Key Areas of Research

Throughout the animal kingdom polarization sensitivity is employed for a broad range of functions (reviews: Wehner, 2001; Labhart, 2016). An understanding of how animals respond to and use the polarization of light is of importance to researchers interested in their sensory ecology. Key areas of research can be summarised under two broad uses of the polarization of light: *contrast vision*, where polarization information is used for object-based visual tasks; and *environmental assessment*, where polarization cues vary over broad spatiotemporal scales and are used to inform navigation or habitat selection behaviour.

#### 1.2.1 Contrast Vision

Contrast vision is an important feature of many visual systems and mediates a wide range of perceptual tasks, such as detecting predators, identifying prey, or visual communication. The role of polarization in these processes is an expanding field of research. One area of focus is the capacity to separate a visual field into objects and background using reflected polarization, roughly analogous to the use of colour vision for image segmentation (*e.g.* How *et al.*, 2015). In colour-sensitive animals, polarization might be used in combination with colour to augment the effectiveness of object detection (*e.g.* Kelber, 1999; Kinoshita *et al.*, 2011), and in colourblind species (such as most cephalopods) polarization may represent a primary means by which image segmentation is achieved (Cronin *et al.*, 2003). Polarization sensitivity might also improve object detection efficiency by permitting the removal of object-concealing scattered spacelight from the visual scene (§2.2.1), enhancing visual contrast (Lythgoe & Hemmings, 1967; Cartron *et al.*, 2013; Sharkey *et al.*, 2015).

Contrast vision is not limited to object detection. There is also evidence that polarization sensitive species of cephalopod mollusc and stomatopod crustacean display polarized body patterns that act as visual signals (Shashar *et al.*, 1996; Cronin *et al.*, 2009; Choiu *et al.*, 2011; How *et al.*, 2014b; Gagnon *et al.*, 2015). In the case of circularly polarized carapace reflections in stomatopods (see §2.2.3), these signals would represent a private communication channel that even other polarization sensitive animals would be blind to (Chiou *et al.*, 2008; Gagnon *et al.*, 2015). It has also been suggested that flowers may signal their profitability to pollinators via patterns in the reflection of polarized light (Foster *et al.*, 2014) and that plant viruses may manipulate polarization reflected from leaf surfaces to attract vectors (Maxwell *et al.*, 2016).

A great range of aquatic insect species, and at least one species of aquatic springtail (Egri *et al.*, 2016), are thought to identify bodies of water from the horizontally polarized light reflected at the water’s surface (Horváth & Kriska, 2008). Horizontally polarized light also triggers oviposition behaviour in swallowtail butterfly *Papilio xuthus*, which uses the angle of polarization of reflected light in combination with colour cues to detect appropriate leaves for oviposition (Kelber, 1999; Kinoshita *et al.* 2011). This is a good example of how polarized light may provide additional information about a visual object once it has been detected.

#### 1.2.2 Environmental assessment

Polarized light may also provide information about more broad-field environmental cues. The best studied example is the incorporation of information from polarized skylight into the celestial compass of many insect species (reviews: Wehner, 2001; Horváth & Varjú, 2004), a capacity that has also been suggested in some crustacean (Bainbridge & Waterman, 1957) and mollusc species (Jander *et al.*, 1963), and even some vertebrates (Taylor & Adler, 1973; Able & Able, 1993; Parkyn *et al.*, 2003). Since the pattern of polarized skylight indicates the sun’s compass bearing (the solar azimuth), it may be used as a reference frame for geographic body-axis orientation when the sun is not visible. A great number of insect species possess a specialised polarization sensitive region in the dorsal eye, the dorsal rim area (DRA), which is used to detect this skylight pattern (Wehner & Strasser, 1985; Labhart & Meyer, 1999). At night, when scattered sunlight is no longer present, moonlight scattered via the same process provides an equivalent polarization pattern that can be used by crepuscular and nocturnal dung beetles, and perhaps other night active insects, for orientation (Dacke *et al.*, 2004; el Jundi *et al.*, 2015).

In aquatic environments polarization may also provide information about water depth. A study involving water flea *Daphnia pulex* found that this species is attracted towards more strongly polarized light (Schwind, 1999), a behaviour suggested to achieve ‘shore flight’ towards deeper water (which often produces more strongly polarized spacelight) and away from shore-dwelling predators. Since polarized spacelight is a feature of many aquatic habitats (§2.2.1), other aquatic species may also use polarized light to guide their shore-flight responses.

While theories on the function of both polarization contrast vision and the use of polarized environmental cues have been under study for some time, at the time of writing both are reaching a new period of improved characterisation and understanding. Recent innovations in the use of Liquid Crystal Displays (see §4.1.3 and §4.2.3) have led to clear demonstrations of the extraordinary polarization vision of cephalopods and crustaceans (Temple *et al.*, 2012; How *et al.*, 2012, 2014; Daly *et al.*, 2016). Careful study of the neuronal processing of skylight polarization cues in insects has led to an improved model for how skylight polarization is interpreted (Pfeiffer & Homberg, 2007; Bech *et al.*, 2014) and combined with other orientation cues (el Jundi *et al.*, 2015). As we come closer to understanding the details of this aspect of the animal visual world that remains alien to our intuition, it continues to be important to focus on the methods we use to produce, control and measure polarized stimuli.

## 2. Polarized Light

### 2.1 What is Polarized Light?

#### 2.1.1 Light—an Electromagnetic Wave

Light is a form of electromagnetic radiation, part of a spectrum that includes X-rays, microwaves and radio waves. In general it passes through space as a *transverse* wave (*i.e.* oscillating at right angles to the direction of travel) that consists of an oscillating electric and magnetic field. For biological systems, we usually only consider the orientation of the electric field, since it is this that directly affects light absorption, and hence vision. Any electromagnetic wave also consists of discrete quantized packets of energy, called photons. Bright light contains more photons than dim light, and the energy of the wave relates to the frequency of the light.

Polarization is, however, a distinct property of light that defines both the relative orientations of the waves as they propagate and how light is reflected, scattered and transmitted through different materials. A beam of light usually consists a large number of waves, and the polarization of a light source concerns the distribution of the orientation of the electric fields of the waves. A beam in which all of the waves oscillate horizontally is called horizontally polarized light (Fig. 2.1A, left). If all of the waves in the beam oscillate vertically then it is vertically polarized (Fig. 2.1A, middle). On average, the predominant axis of the distribution of the waves from a light source is the *angle of polarization* (AoP). The AoP is an angular measure that can vary between 0° and 180°. A second property is the *degree of polarization* (DoP; or percent polarization) and this is the ratio of the (average) intensity of the polarized portion of the beam to its total (average) intensity. For example, unpolarized light, which is composed of multiple waves with a uniform distribution (Fig. 2.1A right) has a DoP of 0, while plane polarized light, in which all waves are oscillating in a single plane, has a DoP of 1. Natural scenes tend to contain light with DoP values ranging between 0 and around 0.5.

The property of *ellipticity* is less straightforward to understand. By describing a wave of light as being composed of two components with perpendicular electric fields, the ellipticity is a measure of the phase relationship and the relative amplitudes of these two components. At any time point, the distribution of the resultant electric field of a beam of light maps out as an ellipse, or in one limiting case, a circle for circularly polarized light (Fig. 2.1C). Most light in nature is not elliptically polarized and the vast majority of animal eyes do not analyse the ellipticity of polarized light (although mantis shrimps provide an exception in both of these cases; see §2.2.3 Biological Polarizers). Because there is no difference between the probability of absorbance of either handedness (rotation direction) of circularly polarized light by visual pigment chromophores, the ellipticity has no effect on the detection of polarized light at the photoreceptor level. This is encapsulated by the measurement of the *degree of linear polarization* (DoLP; see Box 1), which disregards any elliptical component.

**Fig. 2.1.**
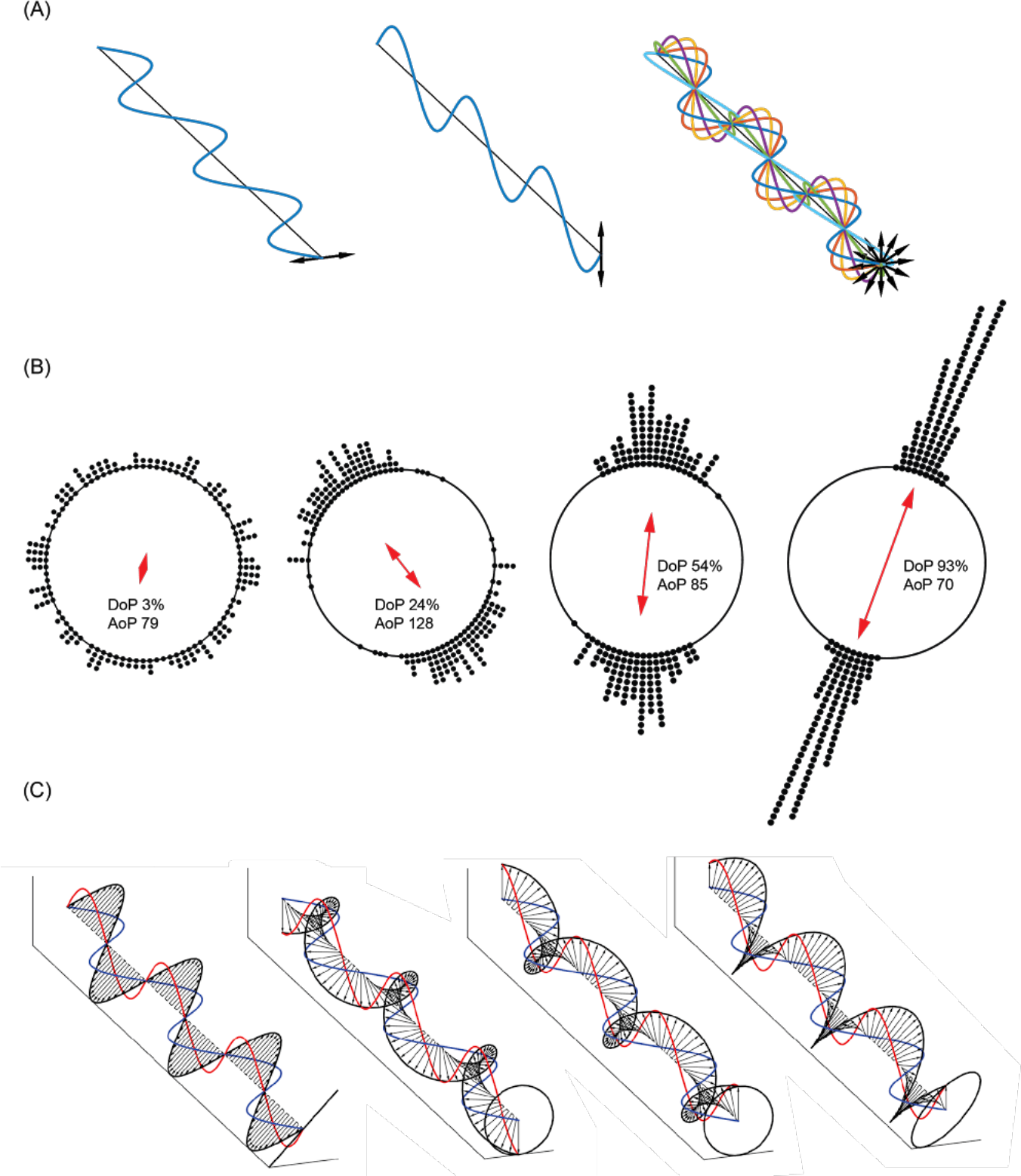
Visualising polarization states.(**A**) Three beams of light propagating toward us along the same axis. Each has a different polarization state. The **left** and **centre** panels show 100% horizontally and vertically polarized light. To the **right** is 0% polarized (unpolarized) light comprising waves that oscillate with a uniform distribution of angles. Shown as a coherent beam to aid visualisation (most light is incoherent: the waves do not have a defined relationship with one another). In (**B**) points on the outside of the circle represent randomly sampled angle distributions of a series of waves comprising a beam, and the arrow within the circle gives the resultant angle of polarization. If the constituent waves oscillate in all directions, the beam is unpolarized, degree of polarization ≈ 0 (**left**). The beam is partially polarized if the distribution of oscillation planes has an overall direction**—**its angle of polarization (**centre**). The degree of linear polarization describes the spread in values (precision of their centre). If all waves oscillate in the same plane, the light is completely linearly polarized: degree of polarization ≈ 1 (**right**). (**C**) Circular polarization and ellipticity, in waves shown as being made up of a vertical (red) and horizontal (blue) component. Ellipticity is governed by the relative phase (distance between peaks) between these two components. A phase difference of zero (or integer multiple of a half wavelength) results in (diagonally) linearly polarized light (**left**). Phase differences of a quarter of a wavelength give left-handed (**left-centre**) or right-handed (**right-centre**) circularly polarized light. Phase differences between these two limits give elliptically-polarized light (**right**). In these example cases, the components’ amplitudes are identical.

###### Box 1: Describing Polarization

**Angle of polarization** (**AoP**; also commonly (but incorrectly) referred to as the e-vector angle or *×*) describes the predominant angle (relative to some external reference *e.g.* horizontal or vertical) along which the electric fields in a light beam oscillate. While this is often referred to as the e-vector angle, the term is not entirely appropriate, since the angle of the electric field vector is a property of single wave and not of a time-averaged beam of light. AoP should not be confused with the polarizing angle, which describes the angle of incidence at which a beam of light becomes polarized after reflection from a surface (see §2.2.2).

**Circular polarization and elliptical polarization** are best described by a wave made up of two electric field components (Fig. 2.1C) directed vertically (red) and horizontally (blue). If their points of maximum and minimum amplitude (peaks and troughs) are aligned (in phase; Fig. 2.1C, left) the resultant electric field oscillation is linear. When these points are out of phase, the resultant electric field traces an ellipse. If this phase difference is a quarter of a wavelength (Π/2) then the electric field vector traces a corkscrew of circular cross-section: circularly polarized light. Any phase difference not an integer multiple of Π/2 (or 0) results in elliptically polarized light. Note however this is only a model to aid our interpretation of observed phenomena.

**Degree of polarization** (**DoP**; also commonly referred to as percent polarization or ∂) refers to the proportion of waves in a source of light that have a particular polarization state. DoP varies between 0 (for unpolarized light) and 1 (for uniformly polarized light). DoP accounts for linear and elliptically polarized light and is calculated from the Stokes parameters (see §2.1.2). If light has an elliptical component, degree of linear polarization (DoLP) may better describe its discriminability for a polarization sensitive animal, since light that is highly linearly polarized or highly elliptically polarized both have high DoP, but the elliptical component must be converted to linear before it can be discriminated.

**Degree of linear polarization** (**DoLP**) describes the proportion of waves in a light source that are oriented in a particular plane. To determine how a light source appears to an animal with linear polarization vision, the DoLP is a practical, relevant measure.

**Optic axis, fast** and **slow axis.** For a material with more than one **refractive index**, the **optic axis** is the axis along which light can propagate, and, irrespective of the angle of polarization, only see (be affected by) one of those refractive indices. The **slow axis** (Fig. 2.1C, red: left centre, blue: right centre) is the orientation for which the polarization is slowed the most by seeing the highest refractive index. Conversely, the **fast axis** is the orientation for which light sees the lower refractive index. Retardation, and the conversion of one form of polarization to another (*e.g.* linear polarized light becoming circular) occurs when the angle of polarization is not coincident with either the fast or slow axis of the material. The resolved components therefore travel at different speeds and a phase difference between the components is introduced.

**Refractive Index i**s the ratio of the speed of light in a vacuum to its speed in a material. The speed of light in many materials depends on the substance’s orientation, and many crystals have two or three different refractive indices depending on a wave’s angle of polarization relative to the crystal axes.

**Transmission axis (TA)**, the axis parallel to the angle of polarization transmitted by a linear polarizer. The ratio of transmission through two polarizers with perpendicular or parallel transmission axes gives their **extinction ratio**.

#### 2.1.2 Stokes Parameters

Stokes parameters are a mathematical representation of the polarization of light and are used to calculate the AoP, the DoP and ellipticity. By setting *I* as the value of the total light intensity and letting *l*_α_ represent the intensity of light that is transmitted through a polarizer with a transmission axis orientated at α, the Stokes parameters (S_0_, S_1_, S_2_, S_3_), can be defined as

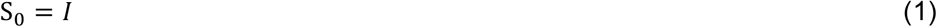

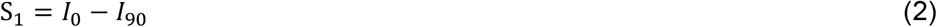

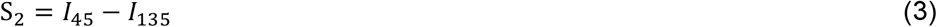

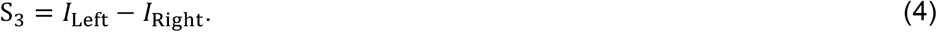

While S_1_ and S_2_ provide information on linear polarization, S_3_ provides a measure of the ellipticity, calculating the difference between the left-handed, *l*_Left_, and right-handed, *l*_Right_, components. Note that S_0_ is the total light intensity.

The angle of polarization, AoP, is given by

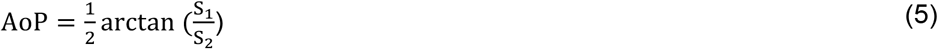

and the degree of polarization, DoP, is given by

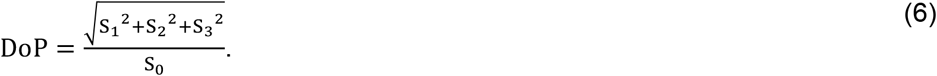

The degree of linear polarization, DoLP, is given by

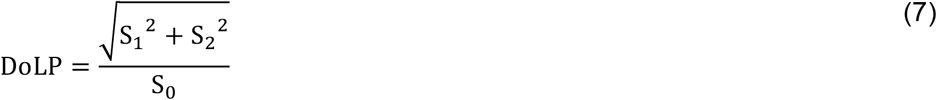

and the ellipticity is given by

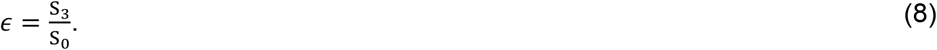

For details of the measurement systems that can be used to calculate the Stokes parameters of stimulus light, see section 3.

### 2.2 Sources of Polarized Light in Nature

Many habitats are rich in polarization information. It is therefore important to consider sources of polarized light in an animal’s natural environment when designing or interpreting a study of polarization sensitivity.

#### 2.2.1 Scattered Light

The most broad-field sources of polarized light in nature are scattered skylight and aquatic spacelight. Through the process of Rayleigh scattering (after: Strutt, 1871) sunlight entering the upper atmosphere is scattered towards a terrestrial observer by molecules smaller than the wavelength of light. The DoP of this scattered skylight increases as a function of the angular deviation from its original path, reaching a maximum at 90°. Additionally, the AoP of this light is at right angles to the initial path, a consequence of the transverse nature of the electric field. Since the angular deviation required for light from the sun to be scattered towards an observer is different in different parts of the sky, the polarization state also differs. The result is a pattern in both DoP and AoP across the sky that can be used, in combination with information about time of day, to determine the sun’s position; a reliable orientation reference when the sun itself is not directly visible (Horváth *et al.*, 2014; Wang *et al.*, 2016). When the moon is the primary source of light within the celestial hemisphere, it too creates a pattern of polarized skylight that can be used as an orientation cue (Gál *et al.*, 2001; Dacke *et al.*, 2004).

In aquatic environments sub-wavelength diameter particles suspended in the water scatter polarized spacelight towards an observer from object-free space within the environment, termed veiling light when it comes between the viewer and a target. Polarized spacelight creates a pattern of polarization, surrounding an underwater viewer, with a high DoP band perpendicular to the predominant direction of downwelling light that tilts with the sun’s elevation (Cronin & Shashar, 2001). It has been proposed (Lythgoe & Hemmings, 1967) that polarization sensitive aquatic species might filter out background spacelight (Johnsen *et al.*, 2011) or veiling light (Cartron *et al.*, 2013; Sharkey *et al.*, 2015) using its polarization, thereby increasing object-background contrast.

Scattered skylight and spacelight represent the most common sources of polarized light in nature; skylight is available to animals in most environments and a variety of aquatic environments permit access to both. The broad-field nature of these cues, and their patterns in angle and degree of polarization, mean that they can be challenging to replicate under laboratory conditions. Often a large sheet of polarizer can suffice in eliciting orientation behaviour similar to that under natural skylight, or simulate a background of polarized spacelight in an aquarium setting, but see sections 4.1.3, 4.1.4, 4.2.1 and 4.2.5 for methods that may be sufficiently versatile to replicate broad-field polarization patterns more precisely, and section 5.1.2 for potential drawbacks of polarizing filters as broad-field stimuli.

#### 2.2.2 Surface Reflections

Specular reflections from most flat surfaces found in the natural world are partially polarized as a function of angle. Particularly prominent are reflections from the surfaces of bodies of water, and as a result many aquatic insects (Horváth & Kriska, 2008) and perhaps some other arthropods (Egri *et al.*, 2016), use polarized reflections to identify their habitats. The leaves and petals of plants (Kelber, 1999; Kinoshita *et al.* 2011) and the exoskeletons of arthropods are other smooth surfaces that produce polarized reflections.

Although most materials found in a laboratory can be used as surfaces that produce polarized reflections, they are not necessarily straightforward to manipulate in terms of their polarizing properties (see §4.1.2). Somewhat counterintuitively, a polarizing filter may function equally well in eliciting polarotactic behaviour in animals searching for bodies of water or smooth leaves, and allow the experimenter to manipulate the angle (*e.g.* Kelber, 1999) or degree of polarization (Egri *et al.*, 2016) without affecting the spectrum or intensity of stimulus light.

#### 2.2.3 Biological Polarizers

In addition to environmentally available polarized light, some species are endowed with structures that control the polarization of light reflected from them (review: Marshall *et al.*, 2014). For instance, many stomatopod crustaceans possess specialised regions in their carapaces that polarize light. Appendages such as the antennal scales and the first pair of maxillipeds make use of the optical properties of highly ordered molecules of astaxanthin and other photonic structures, respectively, to manipulate the degree and angle of polarization of the light that is reflected (Cronin *et al.*, 2009; Chiou *et al.*, 2011, Chiou *et al.*, 2012; Jordan *et al.*, 2016). Additionally, structures that preferentially reflect circularly or elliptically polarized light (see Box 1) have been found on the carapaces of several species of stomatopod, presumably acting as signals for stomatopod species sensitive to circularly-polarized light (Chiou *et al.*, 2008; Gagnon *et al.*, 2015). Polarized reflections may prove to be a widespread feature of stomatopod communication. Many insects also reflect circularly polarized light, however since the discovery by Michelson (1911) over 100 years ago, there still remains very little evidence to suggest any ecological relevance.

**Fig. 2.2.**
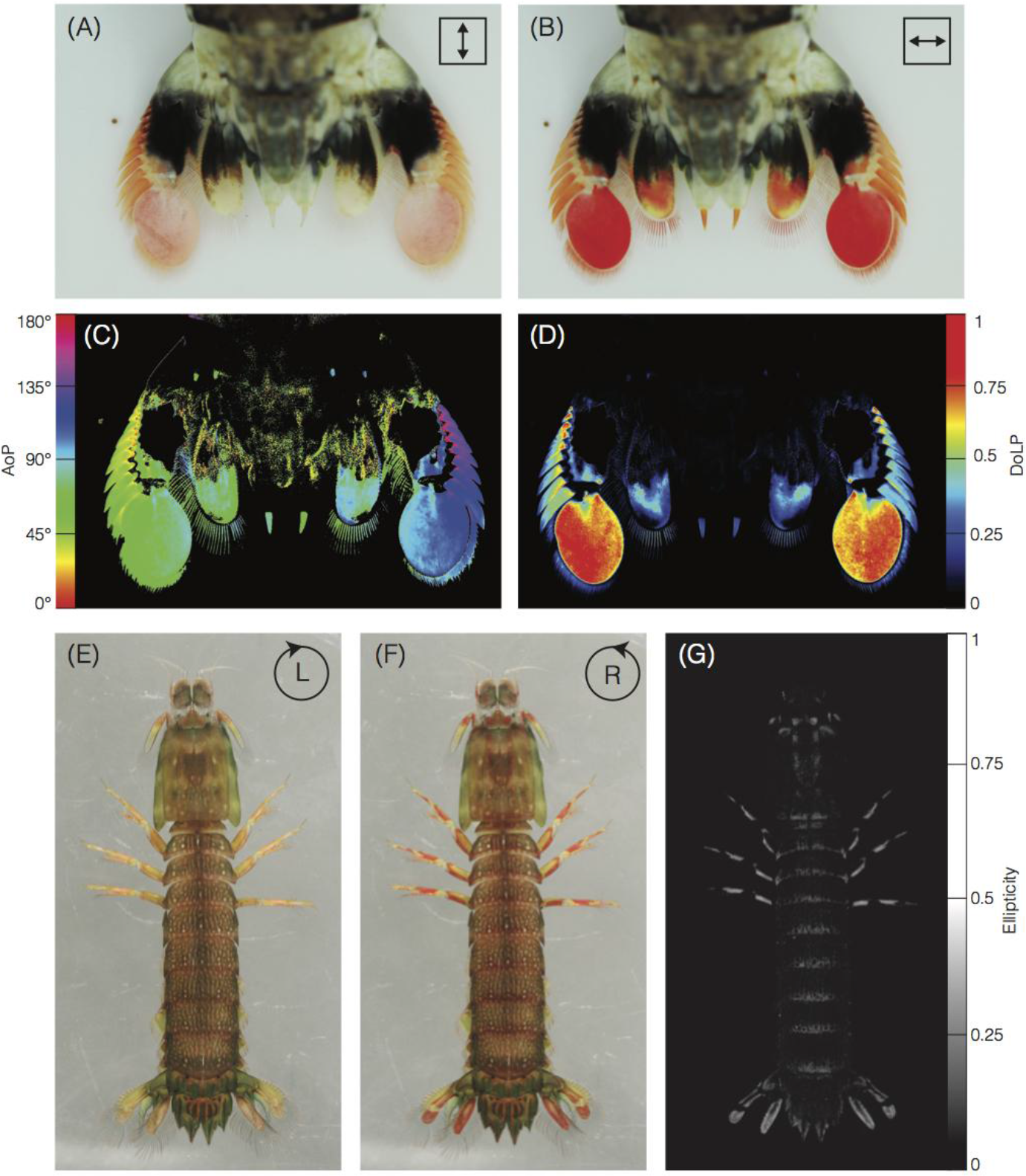
Polarizing structures in stomatopods. (**A-B**) Structures on the telson of *Odontodactylus latirostris* linearly-polarize reflected and transmitted light. Full colour images recorded through a vertically (**A**) and a horizontally oriented (**B**) polarizer, showing the long-pass filtering of horizontally-, but not vertically polarized light (red pigment). (**C**) Angle of polarization as calculated from a digital camera’s green channel (see §3.2.1), indicating that, at these wavelengths, light reflected from and transmitted through the telson is horizontally polarized. (**D**) Degree of linear polarization across the telson, shown as a heat map. (**E-F**) Colour images of *Gonodactylaceus falcatus* recorded through a left-handed and right-handed circular polarizer, highlighting structures on the legs and telson that polarize light with a high degree of ellipticity (**G**).

The iridophores of cephalopod molluscs, such as the cuttlefish *Sepia officinalis* and the squids *Loligo pealei* and *Euprymna scolopes* also reflect light with a higher DoLP than reflections from other parts of the body, forming patterns that can be concealed or revealed via nervous control of the overlying chromatophore layer (Shashar *et al.*, 1996; Shashar & Hanlon, 1996; Chiou *et al.*, 2007; Wardill *et al.*, 2012). Since polarization sensitivity is widespread in cephalopods, it is likely that these too act as polarized signals. Conversely, silvery fishes such as Atlantic herring, *Clupea harengus*, and European pilchard, *Sardina pilchardus*, incorporate a polarization preserving optical mechanism in their skin that, unlike most shiny surfaces in nature (see §2.2.2), does not polarize light via specular reflection at oblique angles. This structure effectively conserves the polarization of an incident beam as it is reflected from the fish’s skin (Jordan *et al.*, 2012; Jordan *et al.*, 2014), helping to better match the intensity of the background illumination.

While visual signals are often highly context dependent, attempts to artificially manipulate (Boal *et al.*, 2004; Gagnon *et al.*, 2015) polarized signals in the laboratory have met with some success, eliciting responses in both cephalopods and stomatopods. Tests of “polarization crypsis”, or lack thereof, in silvery fishes may be carried out via photographic polarimetry (see §3.2.1) or by observing the behaviour of polarization sensitive predators under controlled conditions (*e.g.* Shashar *et al.*, 2000), although neither is straightforward and the application of both methods has met with some criticism (Johnsen *et al.*, 2016).

In general, potential sources of polarization in an animal’s natural environment should be considered when designing experiments in which animals are exposed to polarized light. For a stomatopod that displays green polarized light while defending its burrow, a blue polarized stimulus may be uninteresting. For an insect that orients using polarized skylight, a polarized stimulus that fills only a small portion of its visual field may not elicit orientation behaviour (Henze & Labhart, 2007). To match laboratory stimuli to their presumed natural counterparts, they need to be measured carefully. A comprehensive summary of the best available measurement techniques can be found in the next section.

## 3. How Polarized Light is Measured

As any vision scientist will attest, our own eyes can be unreliable for understanding the properties of light. There are several techniques for measuring the polarization of light, some of which are straightforward enough to be performed in any lab or field setting. Here we describe the basic principles behind these techniques and explain some of the advantages and disadvantages of each approach.

The most common method for measuring polarization requires: a light detector (§3.1.1) and a set of polarization filters (§3.1.2-3; Fig 3.1). The polarization filters convey polarization sensitivity to the light detector, which is otherwise insensitive to the angle or degree of polarization (but see supplement S2). A series of spatially or temporally segregated measurements are then taken through filters of different orientations, and these are fed into simple mathematical formulae to extract meaningful polarization information (§2.1.2).

**Fig. 3.1.**
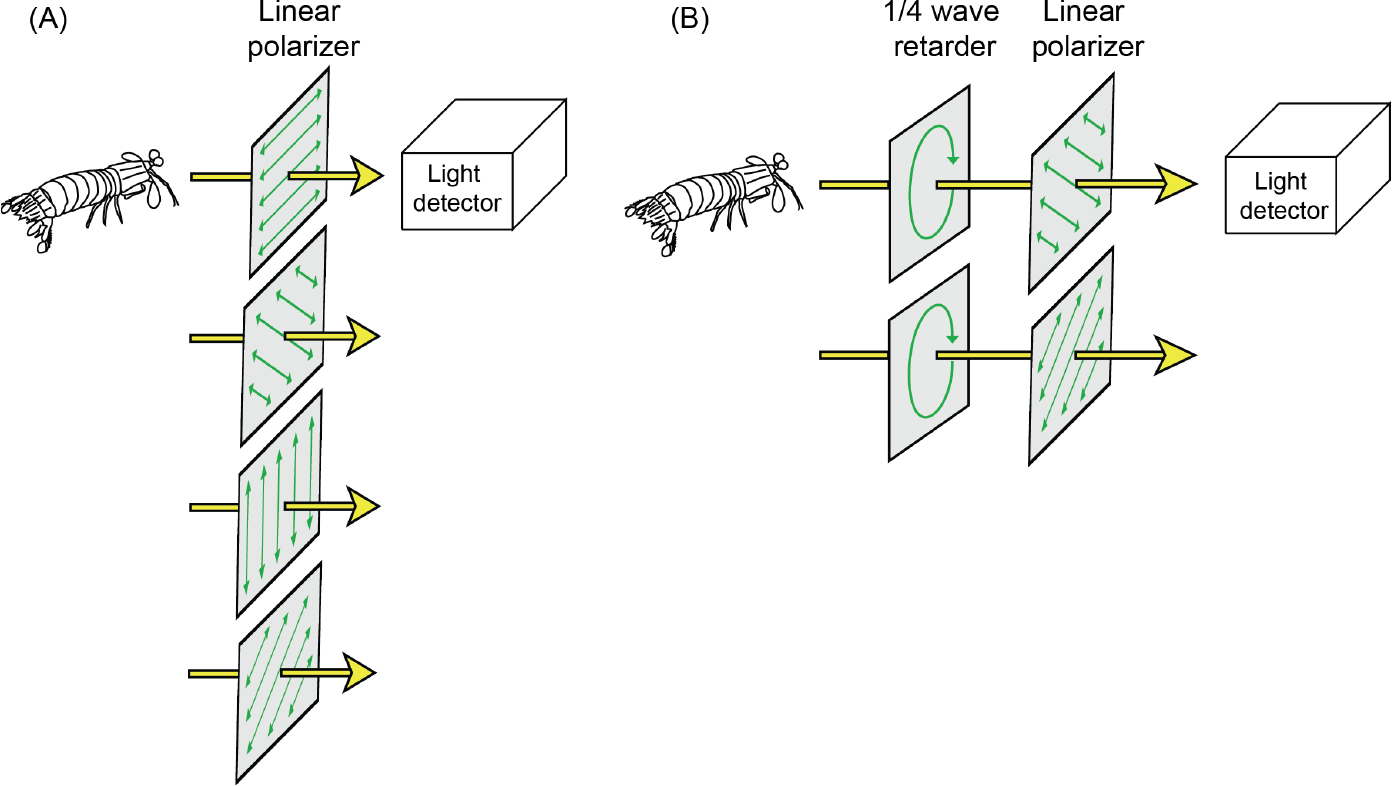
A basic system for measuring the polarization of light. (**A**) For measuring Stokes parameters S_0_-S_2_ the measurements are made with with the polarizers oriented at 0°, 45°, 90° and 135°. (**B**) For measuring Stokes parameter S_3_ a combination of a quarter wave retarder and a polarizer at 45° and 135° is used.

### 3.1 Components of a Measurement System

#### 3.1.1 Light detectors

There are a number of different light detector systems suitable for different kinds of polarization analysis, including photodiodes, photographic cameras and spectrometers, and the choice of approach is generally determined by what information is needed. A detector may be chosen to provide accurate information across the whole spectrum (§3.2.2), or to approximate the visual sensitivity of the animal in question (§3.2.3). In most biological situations, the former is the most desirable method, as few visual systems are well characterised. Furthermore, full spectral measurements of polarization can always be subsequently transformed to model an animal’s spectral sensitivity or used to check for possible artifacts in other parts of the spectral range. However, some situations, such as sky polarization sensors, may benefit from a monochromatic measurement system closer to that employed by animals for the same task, through a reduction in the complexity and variability of the information recorded.

In both cases, there are a number of principles that are important to bear in mind. Firstly, polarization can vary as a function of wavelength, so the spectral sensitivity of the detector is a critical factor. Secondly, most detectors are not linearly sensitive across their stated light intensity range, particularly at the lower and upper limits of this range. This is important given that polarimetry ultimately relies on measuring differences in light intensity (§2.1.2), but can be resolved through calibration. Thirdly, any automatic light adaptation or post-processing mechanisms in the detector must be deactivated to ensure that measurements accurately reflect changes in light levels. Fourthly, it is important to check that none of the optical elements involved in guiding light onto the sensor affect polarization. For example, since diffraction gratings inside spectrometers diffract light differently depending on its polarization state (§S2), light needs to be depolarized before entering the spectrometer, and the same is true of some other optical elements. Lastly, when multiple measurements are taken in series, the device and the light environment to be sampled need to remain as static as possible. Any motion or changing light conditions induce intensity changes unrelated to polarization, resulting in erroneous measurements.

#### 3.1.2 Polarization filters

Because very few man-made light detectors are polarization-sensitive, polarimetry methods rely on a series of polarization filters to differentially absorb specific polarization components. The minimum requirement for measuring the linear component of polarization is a single linear polarization filter, which screens out any light polarized perpendicularly to the transmission axis of the filter, and can range from flexible plastic laminate sheets (§4.1.1), through to high extinction-ratio calcite polarizers, such as Glan-Thompson polarizers.

#### 3.1.3 Retardation Devices

To measure the ellipticity of a polarized stimulus we must add a retardation device. These come in the form of quarter-wave retarders, which are effective at a single wavelength, and Fresnel rhomb retarders, which are effective across a broad spectrum. While quarter-wave retarders convert circular polarization to linear polarization by slowing one component by one quarter of a wavelength relative to the other component (see Box 1), Fresnel rhombs exploit a different optical mechanism and use the phase change in the light that takes place upon reflection.

There are two important factors to bear in mind when choosing the appropriate polarization filters for polarimetry. Firstly, polarization filters and retardation devices are only effective over specific spectral ranges. In particular, many plastic polarizers do not transmit or polarize well in the UV. A different type of polarizer, known as a wire-grid polarizer, is designed for use in the UV, and the calcite Glan-Thompson type polarizers also have better transmission characteristics. Secondly, most of these filters and retarders only perform optimally when the incident light is normal to the surface of the device; off-axis light will usually induce some level of intensity artifact (see §5.1.2); Glan-Thompson polarizers also have a limited acceptance angle. These two factors are relatively straightforward to measure and many manufacturers provide this information with their products.

Here we describe three basic systems that biologists may find useful for measuring polarization. Each has its own advantages and limitations, and each can be modified or enhanced in different ways to improve its effectiveness.

### 3.2 Example Systems

#### 3.2.1 Photographic Polarimetry

The use of photographic cameras for gathering polarization information is one of the simplest and most widely implemented approaches. Because of its simplicity it is also highly prone to misuse and misinterpretation, so an understanding of the benefits and pitfalls of this technique is essential. At its most basic level, all that is required for photographic polarimetry is a standard photographic polarization filter (Fig. 3.2a, i), marked around its perimeter for the desired set of polarizer orientations, and a stable photographic camera with the capacity for full manual control of focus, aperture, exposure and subsequent image processing (Fig. 3.2a, ii). Photographic linear polarization filters and so-called circular polarization (CP) filters are both perfectly adequate for this task, as long as they are mounted on the camera in the standard way. Note that many modern cameras use partially-reflecting mirrors to control autofocus and light metering. These mirrors reflect polarization differently depending on its angle, and incoming light that is highly linearly polarized (as is the case for a linearly polarizing filter) can affect these automatic processes. The problem can be partially mitigated by using CP filters, and by manually controlling focus and exposure.

A typical protocol for photographic polarimetry would be to take a series of images of a scene through the polarization filter angled at 0°, 45°, 90° and 135°, then process this set of images pixel-by-pixel to determine the differences in brightness between them (*i.e.* the component of the scene that is polarized). Converting image contrasts to polarization information can be achieved in several ways (*e.g.* Wolff, 1990; Schechner, 2011; Foster *et al.*, 2014), one of which is based on calculating the Stokes parameters (§2.1.2).

**Fig. 3.2.**
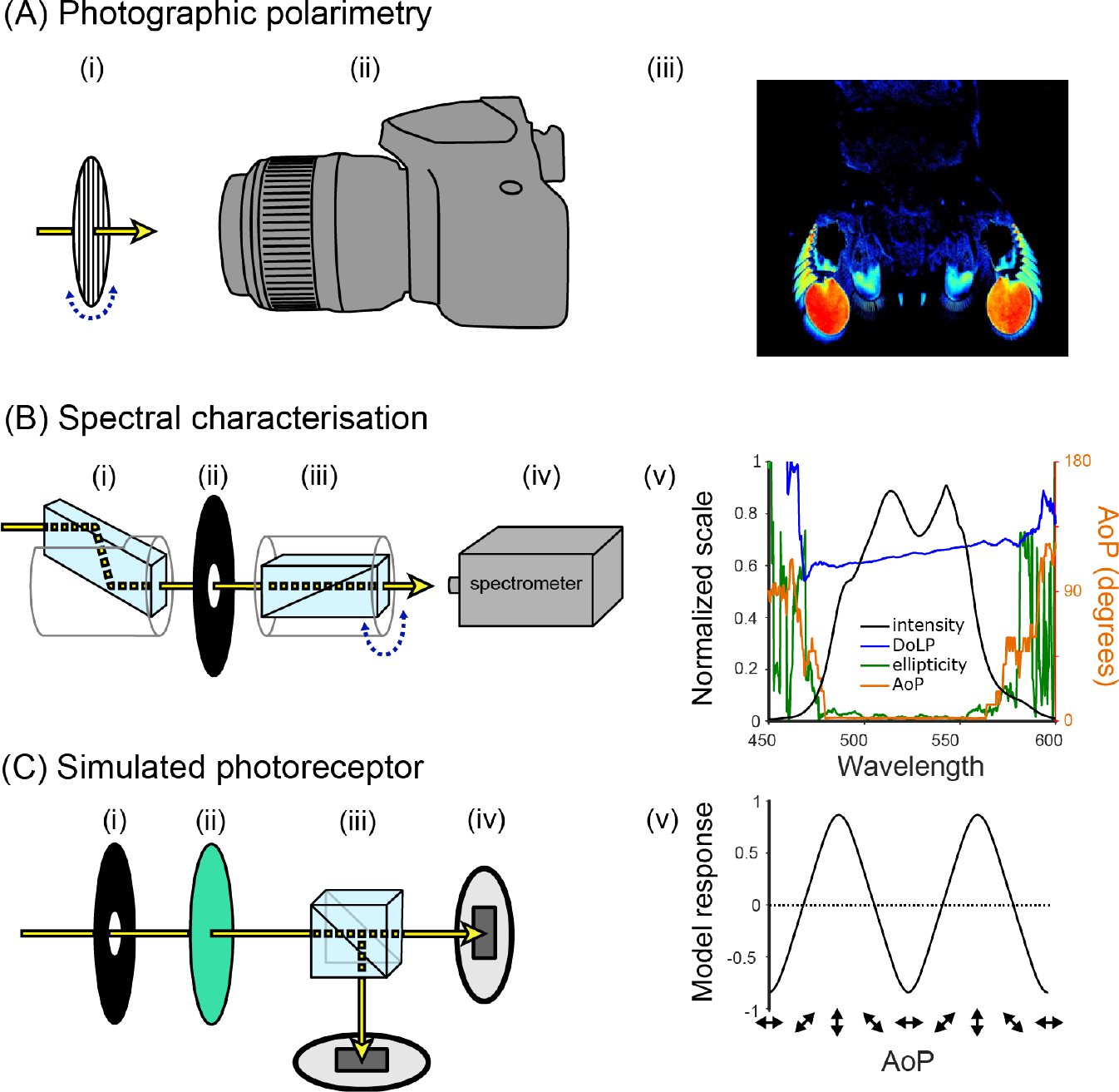
Example systems for measuring polarization and the types of data they produce. (**A**) Photographic polarimetry using (**i**) a rotatable linear polarization filter and (**ii**) a camera in full manual mode. False colour images (**iii**) can be used to represent the spatial distribution of polarization. (**B**) Spectral characterisation of polarization using (**i**) a Fresnel rhomb, (**ii**) an aperture, (**iii**) a rotatable Glan-Thompson polarizer, and (**iv**) a spectrometer. Polarization characteristics may then be plotted against wavelength (**v**). Note: Stokes parameter calculations are only accurate where the intensity is sufficiently high (here: 470-570 nm). The noise at < 470 nm and >570 nm is shown to illustrate the erroneous information resulting from insufficient intensity. (**C**) Simulated photoreceptors using (**i**) an aperture, (**ii**) an interference colour filter, (**iii**) a polarization filter (in this case a polarizing beam splitter), and (**iv**) photodiodes. Output can be simplified into a single measure of contrast between the two detectors (**v**).

Once the polarization information has been calculated, there are a number of methods for presenting it in a visually intuitive manner. This typically involves the use of colour maps (but see recent work by Gagnon & Marshall, 2016, for an alternative) to represent the main aspects of polarization: DoP, AoP, and in some cases, ellipticity (*e.g.* Fig. 2.1). Colour maps may also reflect the contrast sensitivity of the animal, in terms of the threshold at which they can distinguish two different angles or degrees of polarization. This can be achieved by adjusting the bit-depth of the colour map to provide information about what is actually visible to the animal (*e.g.* Temple *et al.*, 2012). It should be noted that no animals are known to perceive the DoP and AoP as separate qualities of light, but it is nonetheless informative to consider spatial patterns of polarization in this way. Some attempts have been made to combine different polarization information within single images, for example by representing the AoP using hue of a colour map and the degree as saturation (Foster *et al.*, 2014), while others have converted polarization information into an estimate of photoreceptor activity in a biological system, which is then represented in false-colour images (How & Marshall, 2014; How *et al.*, 2015).

There are a number of constraints in the technique of photo-polarimetry. Firstly, the camera and the visual scene need to be as stable as possible throughout image acquisition. Edge effects and small movements of the camera between images necessitate an image registration step, and objects moving in the scene can result in false polarization signatures. Secondly, the aperture and exposure controls of the camera must remain unchanged across the image series and be set so that the areas of interest in the scene are neither under nor over exposed. Such situations are often unavoidable in parts of the image (*e.g.* facing into the sun) but should be highlighted as falsely exposed in subsequent processing steps or dealt with using multiple exposures (exposure bracketing). Thirdly, and perhaps most importantly, dark areas of images are particularly vulnerable to producing erroneous polarization signals. This is because, when comparing polarization images in subsequent processing steps, the polarization information is calculated from differences in small numbers (close to the zero end of the sensor scale) that have associated noise. Sensor noise can result in artificially inflated estimates of polarization in these areas (A. Tibbs, I. M. Daly, D. R. Bull and N. W. Roberts, *Bioinspiration & Biomimetics*, under review). We recommend excluding pixels with values in the lower 5% of the sensor range from analysis. Fourthly, care must be taken when using wide-angle lenses with frontally-mounted polarization filters. Towards the periphery of these images light passes through the polarization filter at oblique angles, potentially inducing intensity artifacts (see §5.1.2). Finally, photographic cameras rarely have a linear sensitivity to changes in light levels: *i.e.* contrasts in light levels are measured differently at the bright end of the camera sensor’s range to the dark end. The problem is especially pronounced for film cameras, for which responses to light intensity are non-linear throughout the functional range, but it is also somewhat true of the charge-coupled device (CCD) or complementary metal-oxide-semiconductor (CMOS) chips of digital cameras. This problem can be reduced by deactivating any digital adjustments made to the images by internal processes within the camera (such as auto-white-balance and gamma corrections), and using the raw image format where available (*e.g.* DCRAW: Coffin, 1997). As a final step, the quantitative accuracy of photographic polarimetry measurements can be greatly improved by determining the sensitivity curve of the camera chip (Stevens *et al.*, 2007; J. Smolka and D.-E. Nilsson, in preparation) and lens distortion (J. Smolka and D.-E. Nilsson, in preparation), and compensating for these to linearise the intensity scale.

While there are limitations to the accuracy of photo-polarimetry it remains a useful technique for investigating the relative distribution of polarization information across scenes. It is particularly useful for highlighting areas of interest in natural scenes (such as polarization signals; see §2.2.3), and for discovering possible polarization artifacts in experimental setups (*e.g.* Egri *et al.*, 2016). The recent development of division of focal plane video camera CCD chips, with adjacent pixels etched with different polarization filters may improve temporal, spatial and measuring accuracy in photo-polarimetry, by allowing for the continuous filming of polarization in moving scenes (Powell & Gruev, 2013; York *et al.*, 2014). There have also been developments in engineering camera chips that are inherently sensitive to the polarization of light (Park & Crozier, 2015). Researchers interested in circular polarization patterns can also augment photo-polarimetry using a frontally-mounted quarter-wave retarder filter (*e.g.* Gagnon & Marshall, 2016). Such filters are usually expensive and function only in a restricted part of the spectrum, so care should be taken to analyse only the appropriate colour channel of the camera or introduce a colour filter to restrict the spectral range.

#### 3.2.2 Spectral Characterisation

An effective lab-based system for measuring a single source of polarization involves using a spectrometer coupled to a polarization filter that functions across the light source’s spectrum (Fig 3.2b). Spectrometers are the measurement device of choice among visual biologists. These devices split the measured light across a range of wavelengths using a diffraction grating, and the light is then detected with a CCD chip. Methods for calibrating and using spectrometers are beyond the scope of this review, however, their manufacturers often provide detailed online tutorials (see also: Jonsen, 2012, chapter 9).

The optical properties of the spectrometer’s diffraction grating can induce measurement artifacts if incoming light is polarized. It is therefore important to ensure that the spectrometer receives depolarized light only. While this can be achieved by adding a diffusing component between the polarizer and the spectrometer, the simplest method is to use a long, looped multimode optical fibre, as internal stresses in the coiled optic fibre help depolarize the beam along the fibre’s length (Yu & Rawat, 1992).
Our measurements suggest that a minimum of 2 metres of coiled fibre are required to eliminate this artifact (§S2).

To measure the Stokes parameters, a spectrometer must collect light transmitted through a rotatable linear polarizer. Glan Thompson or Glan Taylor linear polarizers (Fig. 3.2b, iii) are often used, since these polarizers have high extinction ratios across the UV and visible spectrum. To measure the ellipticity of the incoming light, a Fresnel rhomb is added before the linear polarizer in the light path (Fig. 3.2b, i). These devices are intrinsically achromatic (supplement S5), a property that is essential for the characterisation of polarization across a broad range of wavelengths (see supplement S1 for an example setup).

The procedure for measuring Stokes parameters is similar to that used for photo-polarimetry. A series of spectrometer readings are stored for a set of polarizer angles (typically 0°, 45°, 90° and 135°), and then the Stokes parameters S_0_-S_2_ can be calculated for each wavelength (§2.1.2; see also: Wolff *et al.*, 1997). To measure ellipticity, the Fresnel rhomb is added to the light path and two further measurements taken with the linear polarizer rotated −45° and +45° relative to the axes of the Fresnel rhomb.

There are several details to bear in mind when building and using such a device. Firstly, it is important to make sure that the light collected by the spectrometer consists exclusively of light that has passed through the polarization filter(s). The linear polarizer and Fresnel rhomb often come mounted in aluminium holders that do not exclude the possibility of light leaking around its edges. This can easily be remedied by introducing an aperture that restricts the range of angles at which light enters the polarizers (Fig. 3.2b, ii). Secondly, as for the previous example, the measurement apparatus and the light environment need to remain stable during measurement collection. Changes to any of these could create differences between the measurements. Finally, the Fresnel rhomb displaces the optical path of the sampled light (Fig. 3.2b, i), so this needs to be compensated for, especially when measuring small objects.

#### 3.2.3 Simulated Photoreceptors

In situations where the visual system of the study animal is well characterised, measurements can be made to reflect its sensory responses to polarized light more closely. The following example uses pairs of photodiodes to simulate an animal’s polarization-sensitive photoreceptors (Fig. 3.2c, iv). It is important to note the sensitivity, gain, and dark current levels for photodiodes, particularly when comparing input from multiple channels, so a calibration step is likely to be necessary (see Lambrinos *et al.*, 1997, for an example). The measured light beam is filtered by a non-polarizing colour filter, such as a glass interference filter (Fig. 3.2c, ii), to approximate the spectral sensitivity of the animal’s photoreceptor or to restrict the analysis to a specific set of wavelengths. Polarization sensitivity is conveyed by the addition of linear polarizers at angles that match the angular sensitivity of cells in the animal’s visual system (to compare 0° and 90°, a beam splitter might suffice: Fig. 3.2c, iii). As for spectral characterisation, the acceptance angle of each photodiode should be controlled via an aperture (Fig. 3.2c, i). Because the extinction ratios of linear polarizers are far greater than the most effective detectors found in nature, the signal from the photodiode needs to be transformed to better approximate the animal’s polarization sensitivity.

Throughout this manuscript we refer to situations in which measurements of the stimuli and experimental arena are required. Measurements of polarization, radiance and the illumination spectrum help to ensure that the animal observes a stimulus with the intended properties (§4) and is prevented from viewing alternative cues that may confound a study’s conclusions (§5).

## 4. Producing Polarized Stimuli

Just as there are a variety of different sources of polarized light in nature, a range of methods can be used for producing and controlling the polarization of light under laboratory conditions. Although many discoveries have been made in the field of animal polarization sensitivity using nothing more than a sheet of polarizer and an appropriate light source, in recent years a number of more flexible and sophisticated methods have been developed (sections 4.1.3, 4.1.4 and 4.2.3) that permit the production of more complex or naturalistic stimuli. Although these methods place the experimenter in control of the polarization of the visual stimuli, they are not necessarily straightforward or entirely predictable. Where polarized stimuli are used the only sure way to know if their properties meet the requirements of the experiment is to measure them, preferably *in situ,* using one of the methods we have outlined above.

### 4.1 Manipulating Angle of Polarization

While all methods for producing polarized light can be used to manipulate degree of polarization to some extent, the aim, in most cases, is deterministic control of angle of polarization. By far the most common method for doing this is to place a sheet of polarizing filter (§4.1.1) between the light source and the observer. In some instances, specular reflections from smooth surfaces (§4.1.2) have also been used to mimic those from bodies of water. More novel methods, such as the modified LCD monitors (§4.1.3) and “scattering tanks” (§4.1.4) described below, may permit the production of patterns in angle of polarization, adding spatial detail to polarized stimuli.

#### 4.1.1 Sheet Polarizers

Polarizing films, mounted between sheets of plastic or glass, are the most commonly used polarizers in biological studies. Transmitted light is polarized via maximal absorption of a specific angle of polarization coinciding with the direction of the long-axis of absorptive molecules in the film (Land, 1951). The AoP perpendicular to this axis, which is absorbed minimally (and hence transmitted maximally), is termed the polarizer’s *transmission axis* (TA). By rotating the polarizer, stimuli with any AoP can be produced. For this type of polarizer, extinction ratios typically range between 10^−3^−10^−4^. Two main disadvantages of sheet polarizers are spectral transmission and changes in intensity with off-axis transmission (see §5.1.2). Manufacturers typically state the effectiveness of sheet polarizers across the visible range and, although some plastic mounted polarizers perform poorly in the UV, more expensive ones can perform better (supplement S3). Indeed, most studies using sheet polarizers have chosen models, such as Polaroid’s (now out of production) herapathite-neutral ‘B (HNP’B), that transmit and polarize well across the UV-visible spectrum (see §S3).

#### 4.1.2 Specular Reflections

Flat surfaces partially polarize light upon reflection, so the production of polarized reflections requires no specialised equipment. In a series of studies, Horváth and colleagues made use of thin sheets of black polyethylene (review: Horváth & Kriska, 2008) to produce strongly polarized reflections. Panes of glass (Schwind, 1983, 1989, 1991, 1995; Shashar *et al.*, 2005) and oil filled trays (Horváth & Zeil, 1996; Egri *et al.*, 2012) have proved similarly effective. Because these techniques do not require materials that are either expensive or delicate, they are easily applied in field studies, and have led to the identification of more than 250 species of insect as potentially polarotactic.

At a non-metallic surface, a greater proportion of the intensity component that is polarized parallel to a surface (s-polarized) is reflected compared with the component polarized perpendicular to that (p-polarized; *i.e.* polarized parallel to the plane of incidence, the plane containing the incident and and reflected rays). As a result, reflected light is polarized parallel to the reflective surface (Fig. 4.1). The DoP of reflected light varies as a function of the angle of incidence, reaching a maximum at Brewster’s angle, *θ*_*B*_, (Brewster, 1815) with

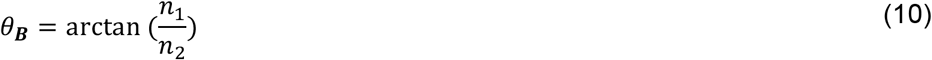

where *n*_*1*_ and *n*_*2*_ are the refractive indices of the initial medium and the reflecting medium respectively. This function limits the range of viewing angles at which high-DoP reflections occur. Since refractive index also varies as a function of wavelength, a polarized stimulus created in this fashion can vary in DoP as a function of wavelength.

**Fig. 4.1.**
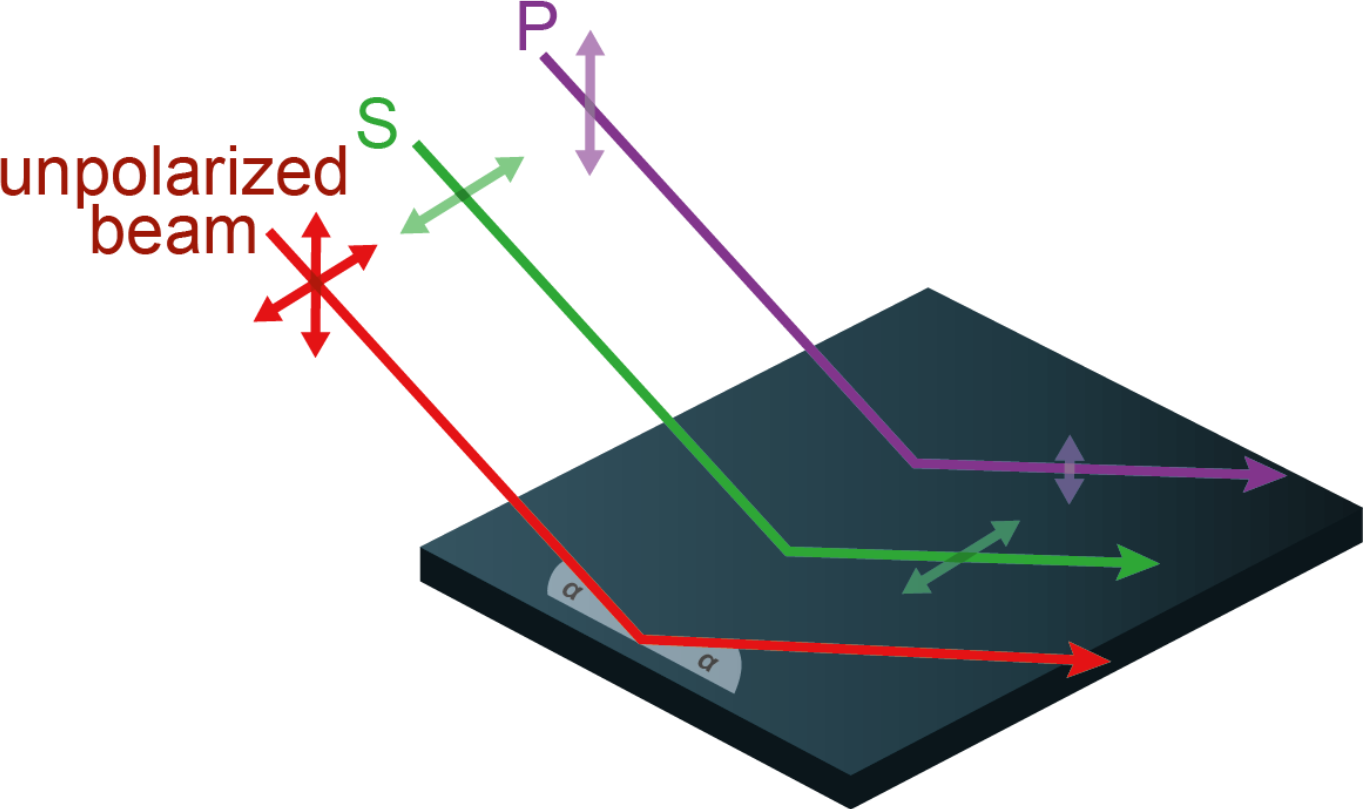
Specular reflection of S-and P-polarization. An unpolarized beam (red) that is specularly reflected from a smooth surface can be considered as a combination of two linearly-polarized components. One of these is polarized parallel to the surface (s-polarized: green) and preserves more of of its intensity on reflection (green double-headed arrow length) than the component polarized perpendicular to it (purple double-arrow length). The angle *α* illustrates that the angle of incidence is equal to the angle of reflection (grey shaded angle).

When reflections are used as a source of polarized light, the spectral and viewing-angle effects require appropriate controls. Control stimuli are typically surfaces with high reflectance averaged across viewing angles, this being a quantity that takes into account contributions from both specular (mirror-like) and non-specular (diffuse) reflection from a surface, including sub-surface scattering and surface reflections from microfacets not aligned with the dominant surface plane (Wolff, 1990). As such, these surfaces are typically lighter coloured equivalents of the same material (*e.g.* Schwind, 1983, 1989, 1991; Kriska *et al.*, 1998; Kriska *et al.*, 2006; Csabai *et al.*, 2006; Horváth *et al.*, 2007; Kriska *et al.*, 2009; Boda *et al.*, 2014). While reflections from these surfaces are lower in DoP, measurements are required to ensure that other cues are not introduced by the change in intensity and reflectance spectrum. For this reason we repeat our earlier recommendation that polarizing filters may be used as an alternative to true surface reflections, where it is specifically the detection of polarization that is of interest.

#### 4.1.3 Twisted Nematic (TN) Liquid Crystal Displays

While polarizing filters are useful for modifying light in a uniform and stable manner, some researchers have turned to liquid crystal display (LCD) technology to produce complex and dynamic polarized stimuli (*e.g.* Glantz & Schroeter, 2006; Temple *et al.*, 2012; How *et al.*, 2012; Daly *et al.*, 2016). These displays, which currently include many flat-panel computer monitors, digital projectors and televisions, generate brightness and colour contrasts by manipulating the polarization of light.

LCDs consist of a liquid crystal panel with electrodes sandwiched between two linear polarization filters oriented at 90° to one another (Fig. 4.2A-B). Twisted Nematic (TN) LCDs can be used to control the AoP of transmitted light. In a TN device the alignment layers are deposited at 90° to each other on the inside of glass of the LCD panel. This induces a 90° twist in the long axes of the liquid crystal molecules (Fig. 4.2A) through the device. This quarter turn of a helix has the appropriate properties to guide the AoP, such that AoP rotates through 90° and the outgoing light is aligned with the transmission axis of the front-most polarizer. This creates a bright pixel. When a voltage above a specific threshold is applied, the liquid crystal molecules reorient, with their long axes becoming parallel to the light path (Fig. 4.2B). In this scenario, light passes through the liquid crystal with its AoP unaltered and is absorbed by the front polarizer. This creates a dark pixel. The addition of red, green and blue coloured filters in neighbouring pixels allows the full control of colour that we are familiar with on our computer monitors.

**Fig. 4.2.**
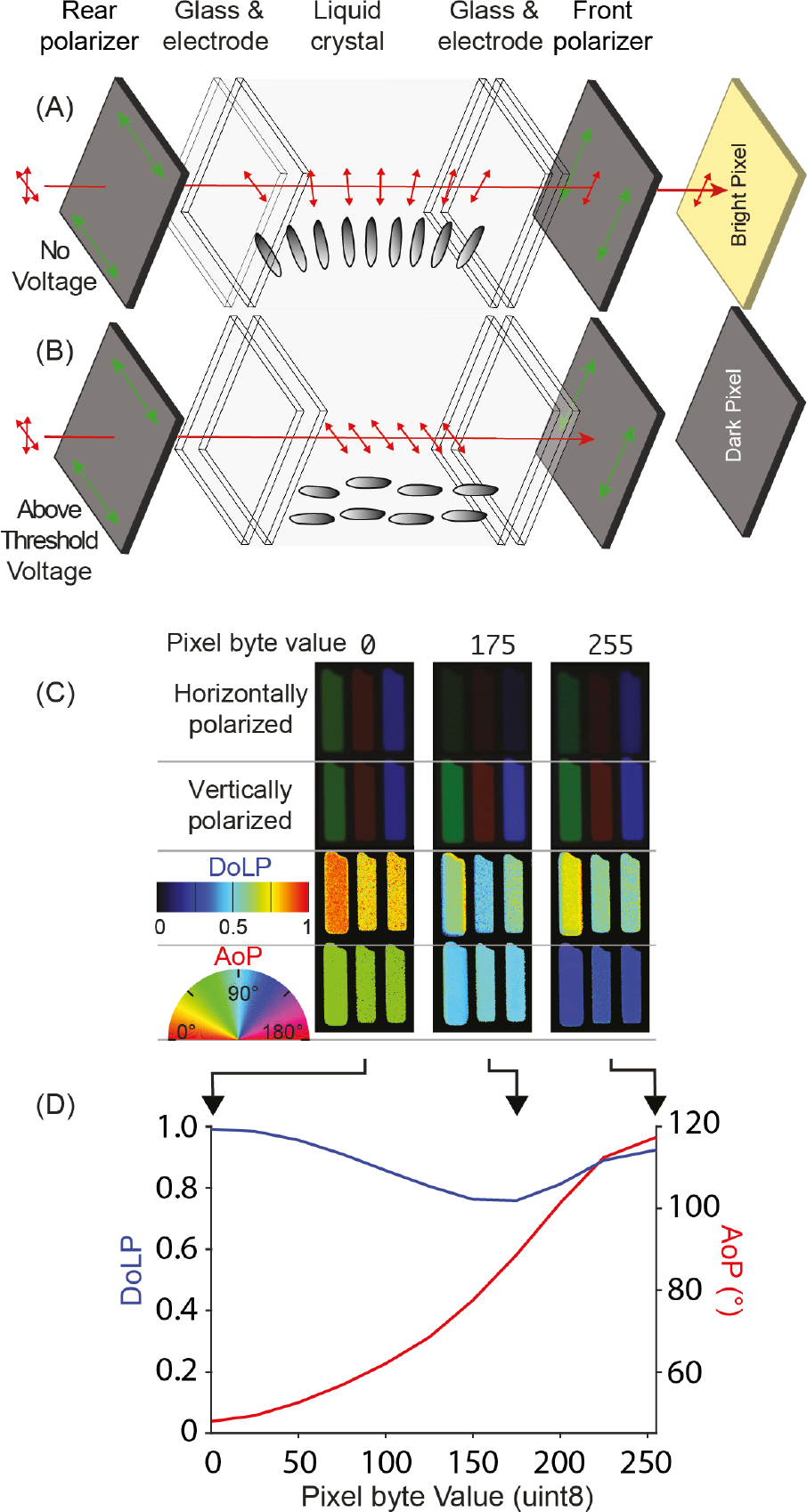
A twisted nematic (TN) panel that can be converted to an angle-of-polarization monitor. (**A**) Schematic of TN LCD Monitor Layers. Illumination (red) from the monitor’s light source first passes through the rear polarizer, for which the transmission axis (green arrows) is perpendicular to the front polarizer. The long axes of the liquid crystal molecules (gray ellipses) twist through 90°, causing a 90° rotation of the AoP. As a result, the light is transmitted through the front polarizer. (**B**) When a voltage is applied, above a threshold, the molecules start to reorient, becoming perpendicular to the glass substrates. At a sufficiently high voltage, the 90° twist is completely removed, and the liquid crystal layer no longer rotates the AoP. As a result, the light is absorbed by the front polarizer. A change between 0° and 90°, via application of an intermediate voltage level, results in a pixel of intermediate brightness. (**C**) Imaging polarimetry (§3.2.1) showing red, green and blue pixels in a TN panel (Dell 1908FPC) converted to an angle-of-polarization monitor via removal of the front polarizer. Pixel byte values for 0 (black), 175 (grey) and 255 (white) shown, each producing a different AoP. (**D**) AoP and DoLP characterisation of the monitor’s output across the 256 interval input scale, showing the gradual change in rotation of the AoP of light as the input value increases (decreases in voltage for a TN panel).

Converting a TN LCD monitor from a colour-and-brightness monitor to a polarization monitor is as simple process and involves only removing the front-most polarizer. The result is that transmitted light is uniform in intensity, but varies in polarization. The exact polarization properties of the transmitted light across the 256 levels of voltage applied to each pixel varies according to: 1) the colour channel; 2) the inbuilt brightness and contrast settings of the monitor; and 3) the brand and type of LCD panel. There may also be measurable variations within models caused by differences in components.

Digital projectors that use a liquid crystal device can also be modified in a similar way. However, the outermost polarizer film can be difficult to access and is sometimes located on three different LCD components, one for each colour channel. Patterned vertical alignment (PVA) LCD monitors can also be used to present polarized stimuli (Fig. 4.4), permitting control over degree-rather than angle of polarization (see §4.2.3).

A number of further modifications can be useful for adapting LCD monitors for polarization experiments. Replacing the inbuilt light source with an alternative, bespoke source can be beneficial for altering the brightness or spectrum of the output light. Opening the rear of the LCD panel to replace the light source usually requires the removal of the power source and control circuitry (which must then be housed separately). Removing unwanted colour channels helps to reduce the spectral complexity of the transmitted light stimulus, simplifying the interpretation of behavioural results. This can be achieved by placing appropriate colour filters between the light source and the rear-most surface of the LCD panel (many colour filters affect polarization and should therefore be placed before the rear polarizer in the light path). Finally, the range of angles of polarization produced by LCD monitors tends to be around 90°, but can be shifted, relative to real-world coordinates, by mounting the monitor on a rotatable stand. For example, a monitor for which AoP can be adjusted between −45° and 45° to vertical, can be rotated by 45° to produce a stimulus that can be varied between 0° and 90°.

The most important tasks throughout all modifications and adjustments are the measurement and calibration of the monitor’s output (see supplement S8 for more detail). Accurate measurements are critical to determine the range of stimuli a monitor produces. LCD monitors seldom produce linear changes in polarization (Fig. 4.2D). For example, for displays with an AoP ranging between −45° and 45°, the change in angle relative to the byte value addressed to each pixel is non-linear and is usually accompanied by changes in ellipticity (and therefore DoLP), in some monitors reaching as high as 60% for intermediate pixel values.

One concern when using LCD monitors for polarization experiments is that they may produce intensity artifacts for oblique viewing angles. Just like polarizing filters (see section 5.1.2), LCDs function optimally for light incident perpendicular to the monitor’s surface (§S7) and as such we recommend that the experimental animal is positioned to avoid oblique viewing angles (§5.2.1). In cases where this cannot be achieved, these potential intensity cues should be measured and taken into consideration.

#### 4.1.4 Manipulating the Angle of Polarization Using Scattering

Although scattering is a common source of polarized light in nature (§2.2.1), under laboratory conditions both the angle and degree (§4.2.1) of polarization can be controlled by using suspended particles as a scattering medium and manipulating the illumination (Fig. 4.3). This method creates a broad-field polarized stimulus, and can be used to mimic veiling light, by combining unpolarized light (transmitted straight through the medium to the observer) with scattered polarization (scattered at 90° towards the observer). An example of a scattering medium might be a mixture of water and subwavelength sized particles, such as those found in skimmed milk (Sharkey *et al.*, 2015), contained within a glass or clear acrylic tank. When the light illuminating the tank from above is unpolarized, a proportion of that light undergoes Rayleigh scattering, creating a weakly polarized light field. If the illumination itself is polarized (*e.g.* by introducing a polarizer before the tank) it follows that a greater proportion of the side-scattered light is polarized.

The AoP of this scattered light can be controlled via the illumination. Since side-scattered light is polarized perpendicular to both its original and final direction of travel (see §2.2.1), the whole arrangement can be rotated around the study animal’s viewing direction, resulting in a light field that is identical relative to the direction of illumination (Fig. 4.3). Rotating the AoP of the illumination by 90° also results in a shift in the predominant AoP of the light scattered within the tank by 90° for the original viewing direction. It should be noted that this also reduces the intensity of the scattered light in this direction, which can be compensated for by brightening the illumination. This also reduces the degree of polarization of light scattered in the original direction, which cannot be easily compensated for, but has applications in itself (see below). These techniques are intended to mimic scattered light in an aquatic environment, which may conceal objects (Lythgoe & Hemmings, 1967) or form a polarized background from which objects must be distinguished (Jonsen *et al.*, 2011). One application is to test the advantages of polarization sensitivity in contrast sensitivity through veiling light, using a naturalistic polarized light field (Sharkey *et al.*, 2015).

**Fig. 4.3.**
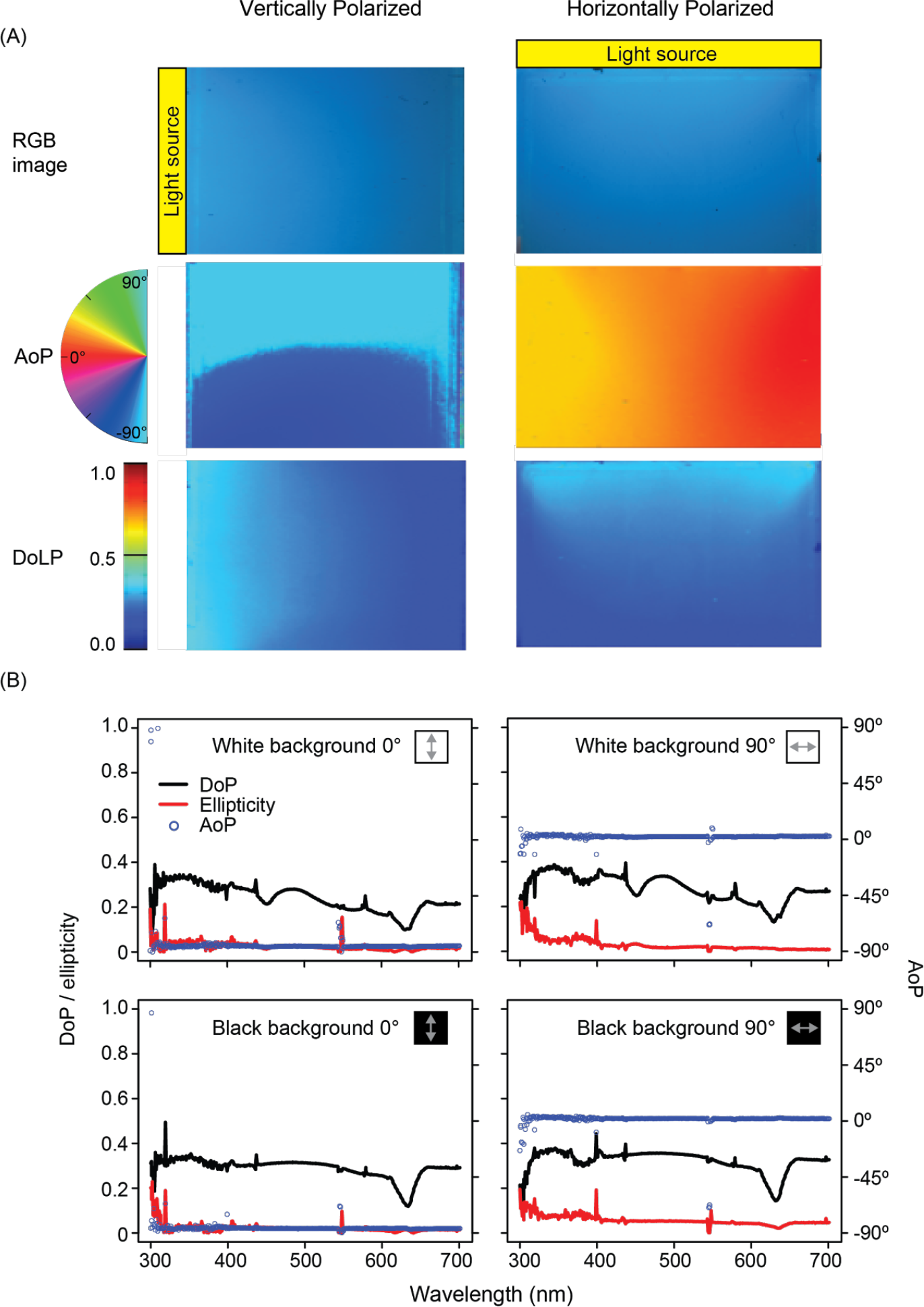
Scattering tank used to manipulate AoP. (**A**) Light source and polarizer either to the side of the tank (**left**) or on top of the tank (**right**) producing vertically-and horizontally-polarized scatter respectively. Colour images (**top**), photo-polarimetric images of angle of polarization (**middle**), and degree of linear polarization (**bottom**). (**B**) Spectral polarization measurements, with a monitor displaying either white (**top**) or black (**bottom**) transmitted behind the tank. *N.B.* At the monitor’s emission peaks (450 nm, 550 nm, 620 nm) the white unpolarized background reduces the observed DoP.

### 4.2 Manipulating Degree of Polarization

Most experiments testing animal polarization sensitivity to date have used polarizing filters (see §4.1.1) that produce polarized light with a DoP approaching 100% but, under natural conditions the degree of polarization is often lower: celestial polarization patterns rarely exceed 85% (Cronin *et al.*, 2006; Horváth *et al.*, 2014) and underwater polarization typically only exists to a maximum of 50% (Lerner, 2014). To assess an animal’s response to polarized light under more natural conditions, stimuli should be produced with DoP values comparable to those in its natural visual environment. Such stimuli also make it possible to characterise the limits of the degree of polarization that an animal can detect, *i.e.* measure the DoP threshold (Labhart, 1996; Glantz & Schroeter, 2006). Methods for controlling the DoP fall into four categories: scattering/diffusion (§4.2.1), adding unpolarized light (§4.2.2), modified LCDs (§4.2.3), and retardation devices (§4.2.4).

#### 4.2.1 Manipulating Degree of Polarization Using Scattering

Scattering can be a source of polarized light (§2.2.1), but it can also decrease the DoP (*e.g*. clouds). The difference is determined by the type of scattering, which depends on the size of the scattering particles and their density. If the particles are very small (*i.e.* less than λ/15, where λ is the wavelength of light) then Rayleigh scattering occurs (see §2.2.2). If the particles are larger (*i.e.* approaching the size of λ) then Mie scattering occurs (Mie, 1908). Mie scattering and multiple Rayleigh scattering events can be used to alter the effective DoP of a transmitted beam. In the laboratory, suitable methods to scatter light involve particulates (*e.g.* sand or hollow glass spheres) suspended in liquids (see §4.1.4) and translucent materials.

A simple scattering filter can be constructed by using the scattering tank method describe above, and keeping it well mixed. The DoP can then be altered by varying the concentration of scatterers or the path length of light passing through the mixture or the size of the particles. Key considerations are: that the tank and scatterers do not vary the spectral characteristics between different DoP conditions and that the tank does not vary the ellipticity of the light. If the specific density of the particulate differs from that of the liquid, it may settle at corners and edges of the tank and it is therefore important to measure the system after it has been set-up and mixed for some time.

Translucent materials offer the easiest means of decreasing DoP, and commercially available diffusion filters can be obtained from photographic and theatrical suppliers, as well as makers of optical components. These are glass, polymer or thin film (gel) diffusers with varying levels of scattering ability. They are not sold as DoP filters nor do they come with any quantification of how they affect DoP, but are nonetheless effective. High-end diffusers may provide a point spread function, describing the distribution of diffused light, and lower-end diffusers may give an arbitrary value that indicates the extent of scattering. The interaction of diffusers with polarization depends, in part, on the distance of the diffuser from other components in the light path: a diffuser positioned close to a polarizer has less of an effect on DoP than one further away.

Among high-end diffusers, frosted quartz best transmits UV, with frosted glass or plastic diffusers only effective at depolarizing light between 400-800 nm. In addition, glass and quartz diffusers do not induce elliptical polarization; as opposed to frosted plastics, which can induce ellipticity when heat stressed. However, glass and quartz can be prohibitively expensive, especially for large filters. Alternatively, purpose-made frosted diffusers can be custom made via sandblasting or acid etching. Numerous filters can also be stacked to decrease the DoP (*e.g.* Egri *et al.*, 2016), but note that the effects of multiple filters are not, in general, linearly additive, and so each condition produced this way must be measured separately. Frosted film (polyester/gel) diffusion filters are available in a wide range of densities, as are grid cloths and spun materials. These are cheaper than frosted glass, plastic or quartz and may be preferable for larger stimuli. One important caveat when using thin film diffusers to control polarization is that these films can act as retarders. As a consequence, the orientation of the diffusers must be controlled and maintained relative to the polarizer to control the DoP consistently.

Various researchers have used scattering to decrease the DoP in their experiments. Hawryshyn and Bolger (1990) used latex microbeads of 1 μm diameter to decrease the polarization of stimulus light. Henze and Labhart (2007) used 2 sheets of translucent tracing paper on a sheet of polarizer to reduce DoP to zero, reversing the order to get a DoP of one. Cartron and colleagues (2013) used “fine natural sand” of <1 mm diameter to decrease DoP and intensity contrast of images displayed on a modified LCD and CRT monitor respectively. Temple and colleagues (2015) used “custom-made volume diffusers” to decrease the DoP of stimuli, based on the same principle of adding particulates to a transparent medium. In each case, since the process of depolarization in a scattering medium is not easy to predict, the measurements of degree of polarization were reported alongside estimates of the DoP or target detection thresholds, as evidence that the scattering media reduced the DoP of stimulus light.

#### 4.2.2 Adding Unpolarized Light

Another method to decrease the DoP of an already polarized light source or field, is to add unpolarized light into the optical path, reducing the proportion of light that is polarized when it reaches the viewer. A simple mechanism for doing this is to position a sheet of glass at an angle between the viewer and the polarized light field. An unpolarized light field can then be reflected from the glass into the optical path, and by adjusting the relative intensity of the polarized and unpolarized light fields, a range of DoP values can be attained. A drawback of this system is that the viewing angle for which it is effective is limited. While the authors have used this approach in a preliminary experiment, we know of no published work that has employed the beam splitting approach in animal behavioural studies.

#### 4.2.3 Patterned Vertical Alignment (PVA) Liquid Crystal Displays

As described earlier (§4.1.3), LCD monitors can be modified to present dynamic or static polarization contrasts instead of intensity and colour contrasts. While models of twisted nematic (TN) LCDs can be modified to produce changes in the AoP across the screen (Glantz & Schroeter 2006; How *et al.*, 2012; Temple *et al.*, 2012; Daly *et al.*, 2016), other LCD technologies can also change the DoP. The differences between LCDs that produce changes in the AoP *versus* the DoP relate to the alignment of the liquid crystal. In patterned vertical alignment (PVA) LCDs, the liquid crystal molecules are oriented parallel to the light path (a homeotropic orientation) in the default state when no voltage is applied (Fig. 4.4A-C). As a result, the polarization of transmitted light is not modified by the liquid crystal, producing a dark pixel.

In a PVA monitor, the electrodes are structured such that they are laterally offset between the two substrates. In addition, the electrodes are also shaped into chevrons, which together with the offset, creates four different domains per pixel when a voltage is applied. The molecules reorientate into the plane of the device at angles of 45°, 135°, 225° and 315° relative to the rear and front polarizers transmission axes. The change in the liquid crystal orientation causes a retardation effect, with different applied voltages creating different ellipticities. However, the symmetry of the four elliptical domains means the transmitted light is always comprised of equal quantities of lefthanded and right-handed polarization. These cancel each other out, resulting in a changing DoP. The PVA monitor that has been used in all experiments to date, (1905FP, Dell, Round Rock, USA) can vary the DoP from 0 to 1, which allows for testing of the minimum difference in DoP between a stimulus and its background that an animal can detect (How *et al.*, 2014). The monitor can also be rotated, to determine if this threshold changes at different orientations relative to the animal’s eye (How *et al.*, 2014).

**Fig. 4.4.**
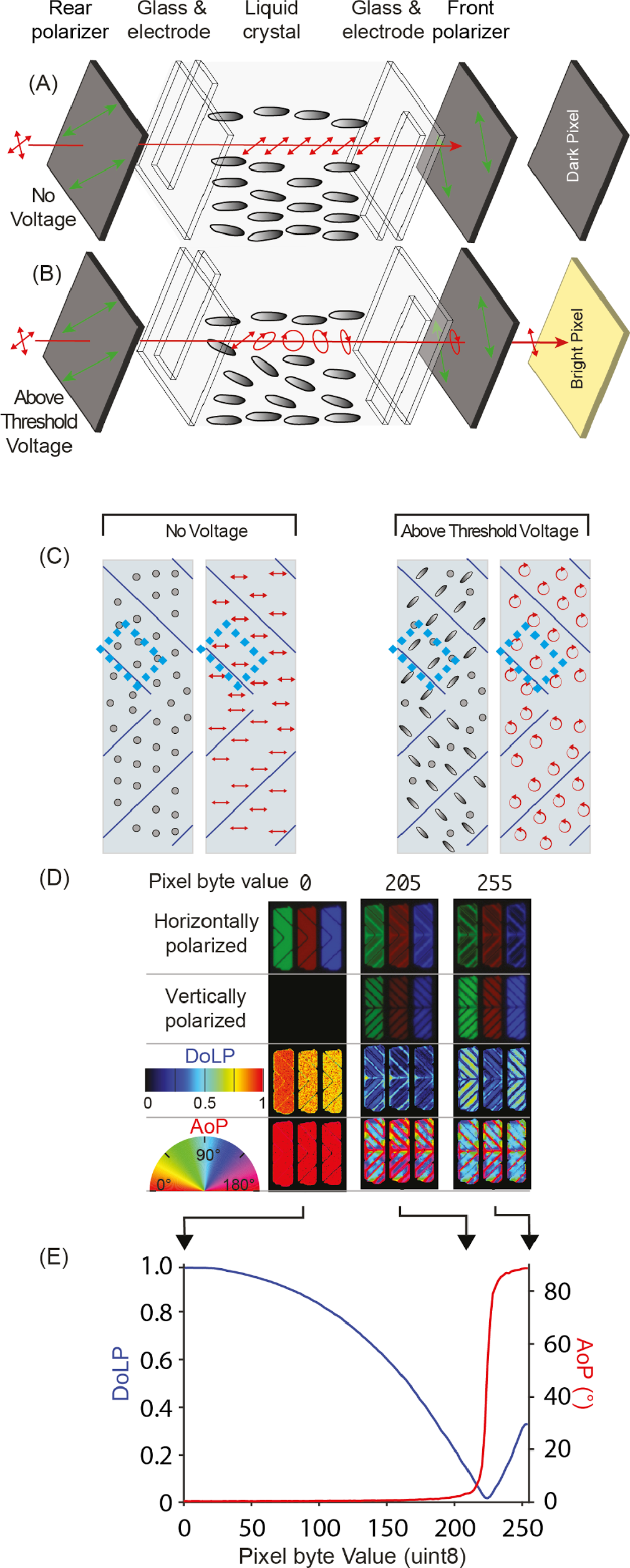
A patterned vertical alignment (PVA) panel that can be modified to act as a degree-of-polarization monitor. (**A**) Schematic of PVA panel layers. The transmission axes of the rear and front polarizer are perpendicular, and a dark pixel is achieved by allowing the illumination (red) to pass through the liquid crystal unmodified. (**B**) For different applied voltages, the liquid crystal layer retards output light to different extents, altering its DoLP. (**C**) Front view of a pixel, showing the liquid crystal alignment (grey ellipses) and polarization state of output light (red arrows). The blue boxes indicate the area of the pixel represented in (A) and (B). The two diagrams show the cases of no voltage (**left**) and an above threshold voltage (**right**). For an above threshold voltage, linearly polarized light is converted to equal quantities of left and righthanded elliptically polarized light in the different domains of the chevron, which combine to cancel out, and change the DoP of the output light. At a particular voltage, the domains act as quarter-wave retarders, changing the polarization into left and righthanded circular polarization, which cancel to give a DoP of zero. (**D**) A modified DoLP monitor (Dell 1905FP) at the pixel level. Polarization changes in PVA displays are complex, varying within domains in each pixel. Conversion of horizontally to vertically-polarized light begins at intermediate byte values (*e.g.* 205), in the regions of the pixel domains in which the liquid crystal molecules tilt. (**E**.) Across most of the input range, the averaged AoP of the monitor’s output remains the same while DoLP decreases with increments in pixel byte value (*i.e.* increases in voltage for a PVA panel).

#### 4.2.4 Retarders

If the study animal is not differentially sensitive to the ellipticity of polarized light, a stimulus’ DoLP can be adjusted by increasing its ellipticity. When the AoP of the linearly polarized light beam entering a quarter-wave retarder is at 45° to the fast and slow axes of the retarder, it converts linear polarization (ellipticity = 0, DoP = 1, DoLP = 1) into circular polarization (ellipticity = 1, DoP = 1, DoLP = 0). If the AoP of the input light is aligned with either the fast or slow axis of the retarder, then there is no effect on polarization and it remains linearly polarized (ellipticity = 0, DoP = 1, DoLP = 1). If the AoP of the input light is at any other angle to the fast or slow axis, the the light is transmitted as elliptically polarized light (*i.e.* 0 < ellipticity < 1, DoP = 1, 1 > DoLP > 0). It should be noted that the value of retardation is not just a function of a material’s optical properties and thickness alone, it is also a function of wavelength (see supplement S4). Some materials act as retarders even though this is not their primary function, for example thin film diffusers and coloured theatrical gels made out of thin polyester or polycarbonate. When using such coloured and scattering filters, care should be taken to place them between the light source and polarizer (rather than between polarizer and animal) and measure any effects of orientation on polarization.

Retarders have been frequently used for decreasing DoLP in behavioural experiments. Henze and Labhart (2007) used overhead transparency film and rotated the fast axis relative to the polarizer to vary DoLP from 1.0 to 0.0 in their experiments with field crickets. How *et al.* (2015) recently used a polymer film quarter-wave retarder (#88-252, Edmund Optics) when investigating the use of polarization contrast by fiddler crabs. Other retarders used include: drafting film (Mylar Medium Weight 0.003, Graffix: Tuthill & Johnsen, 2006), photocopier transparency (Pfeiffer *et al.*, 2011), polycarbonate gel filter (clear #00, Roscolux: Shashar *et al.*, 2000) and laboratory film (Parafilm^®^, Bemis Flexible Packaging: Glantz & Schroeter, 2006).

One of the main caveats in using retarders is that most (including optical-grade zero-order and multi-order retarders) alter the polarization as a function of wavelength, and are often effective over a fairly narrow spectral range (§S4). Thus, the wavelength of stimulus light should be restricted, or the spectral sensitivity of the polarization photoreceptors must be known to fall within the desired range. As for all other methods for the production of polarized stimuli, careful measurement with reference to any information available on the animal’s visual system is vital for control over experimental conditions.

#### 4.2.5 Projected Polarization

While this review was being compiled, a new method was published for controlling both angle and degree of linear polarization in projected polarized stimuli (Stewart *et al.*, 2017). Taking inspiration from a study in which light from a TN LCD was projected onto a translucent screen, producing moving stimuli that contrasted in AoP (Glantz & Schroeter, 2006), the authors modified a digital light processing (DLP) projector to project moving patterns of polarized light (Stewart *et al.*, 2017). While the LCD-based system produced stimuli with one DoLP level at a given time (using a laboratory-film retarder), the DLP-based system produces projected dynamic patterns that can be varied in AoP, DoLP, intensity and even wavelength (although this full range was not employed experimentally).

## 5. Confounding Cues in Experiments

In experiments with polarized stimuli it is important to control for confounding sources of stimulation. Since many of the optical effects that are responsible for the polarization of light also affect its intensity and spectrum, it is always necessary to control for such changes where they may co-occur. These effects can be subtle and in cases where stimuli include UV light these confounds may not be detectable to the human eye. Failure to control for confounding cues can allow an animal’s response to be interpreted as polarization sensitivity incorrectly or, conversely, can distract from true polarization cues. In this section, we provide details of some potential sources of confounding cues in polarized stimuli (§5.1) and describe some methods to control for them (§5.2).

### 5.1 Sources of cue/confounds

#### 5.1.1 Surface Reflections

Specular reflections are a common and recurring issue in experiments with polarized stimuli (discussed in: Taylor & Adler 1973; Coemans *et al.* 1990; Goddard & Forward 1991; Muheim 2011). Because a greater proportion of light that is polarized parallel to a surface (s-polarized) is reflected, compared with light that is polarized in the plane of incidence (p-polarized), the reflected intensity of a linearly-polarized beam changes as a function of the AoP (Fresnel, 1823). This causes the surface to appear lighter or darker depending on the AoP of the illumination. Light that is not specularly reflected or absorbed by a surface contributes to its diffuse reflectance (see §4.1.2), which affects a surface’s general lightness of appearance. While dark surfaces with low diffuse reflectances have been used to minimise the intensity of reflections in a number of studies of polarization sensitivity (Rossel *et al.*, 1978; Mussi *et al.*, 2005; Sakura *et al.*, 2012), this decrease in average reflectance increases the contrast between reflected s-polarized and p-polarized beams (Fig. 5.1), making the stimulus’ AoP discriminable via the brightness of specular reflections. This occurs because a smaller proportion of the beam’s intensity is reflected diffusely, irrespective of the AoP, but the same proportion is reflected specularly, as a function of the AoP.

**Fig. 5.1.**
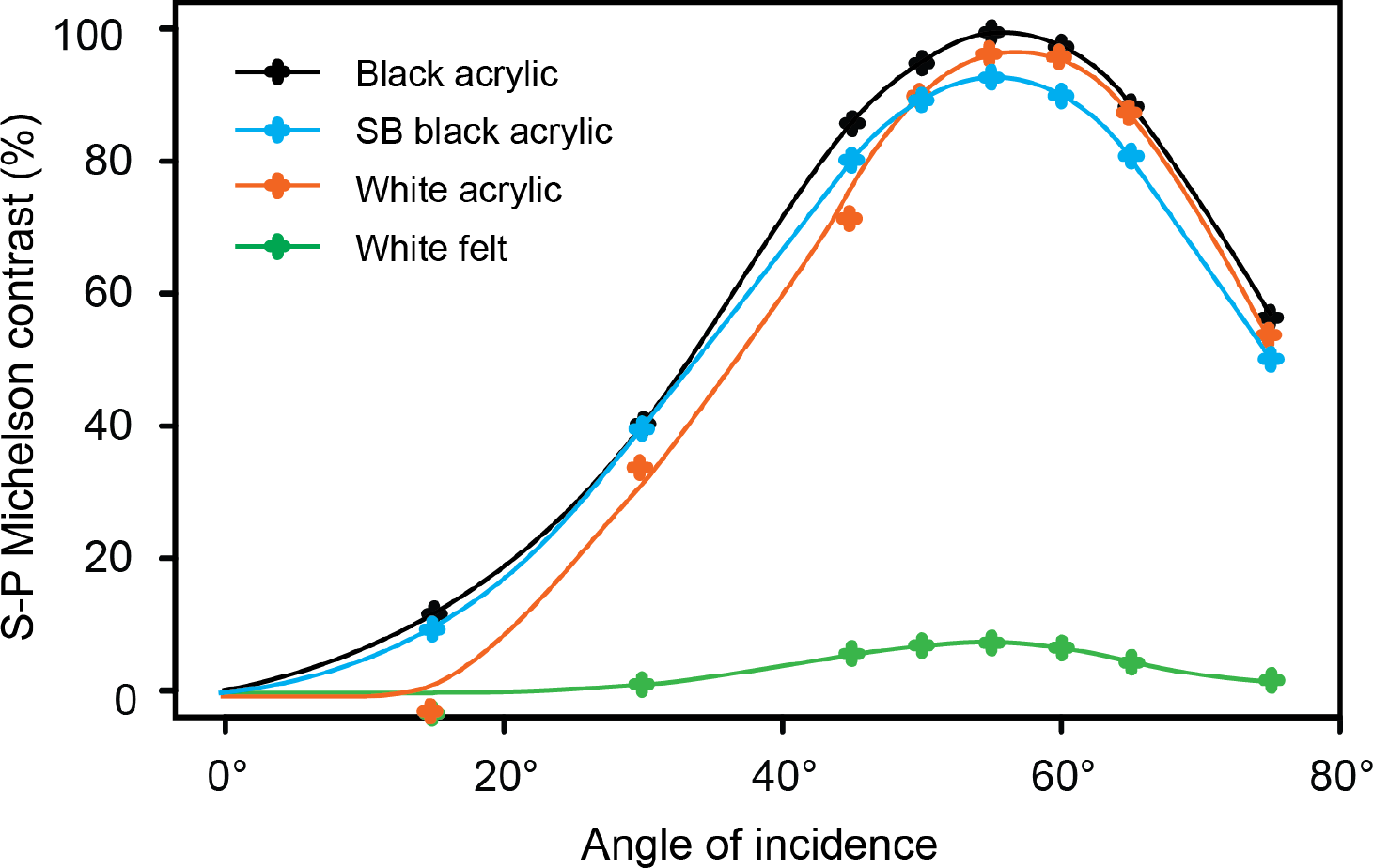
Contrast between reflected p-and s-polarized light. (**Black**) black acrylic (Perspex, Weybridge, UK). For angles of incidence 45-60° almost none of the p-polarized beam’s intensity is reflected light. (**Blue**) white acrylic (Perspex, UK). The difference in reflected intensity between s-polarized and p-polarized light is lower than for black acrylic. (**Orange**) the same block of black acrylic as above, sandblasted to create a ‘rough’ surface. The increase in contrast with angle of incidence is more gradual than for untreated black acrylic. (**Green**) white felt (Fabric Land Ltd., UK). This material is both highly reflective and ‘rough’ (fibrous), and hence the contrast between reflected s-polarization and p-polarization is low at all angles. Raw measurements of reflected intensity available in the supplement (§S5).

Some more recent studies have controlled for reflected intensity cues by using surfaces that are matt, ‘rough’, or highly reflective (e.g. Mäthger *et al.* 2011; Calabrese *et al.* 2014; Melgar *et al.* 2015; Egri *et al.*, 2016). These types of surface can help minimise the difference in intensity between s-and p-polarized reflections as a proportion of overall reflected intensity. Fibrous materials, such as felt and other fabrics, reflect and scatter incident light at a wide range of surface angles (Fig. 5.2). These two effects may be combined by using light-coloured fabric surfaces, with both a high diffuse reflectance and a high proportion of fibres oriented perpendicularly to the surface (Fig. 5.2).

**Fig. 5.2.**
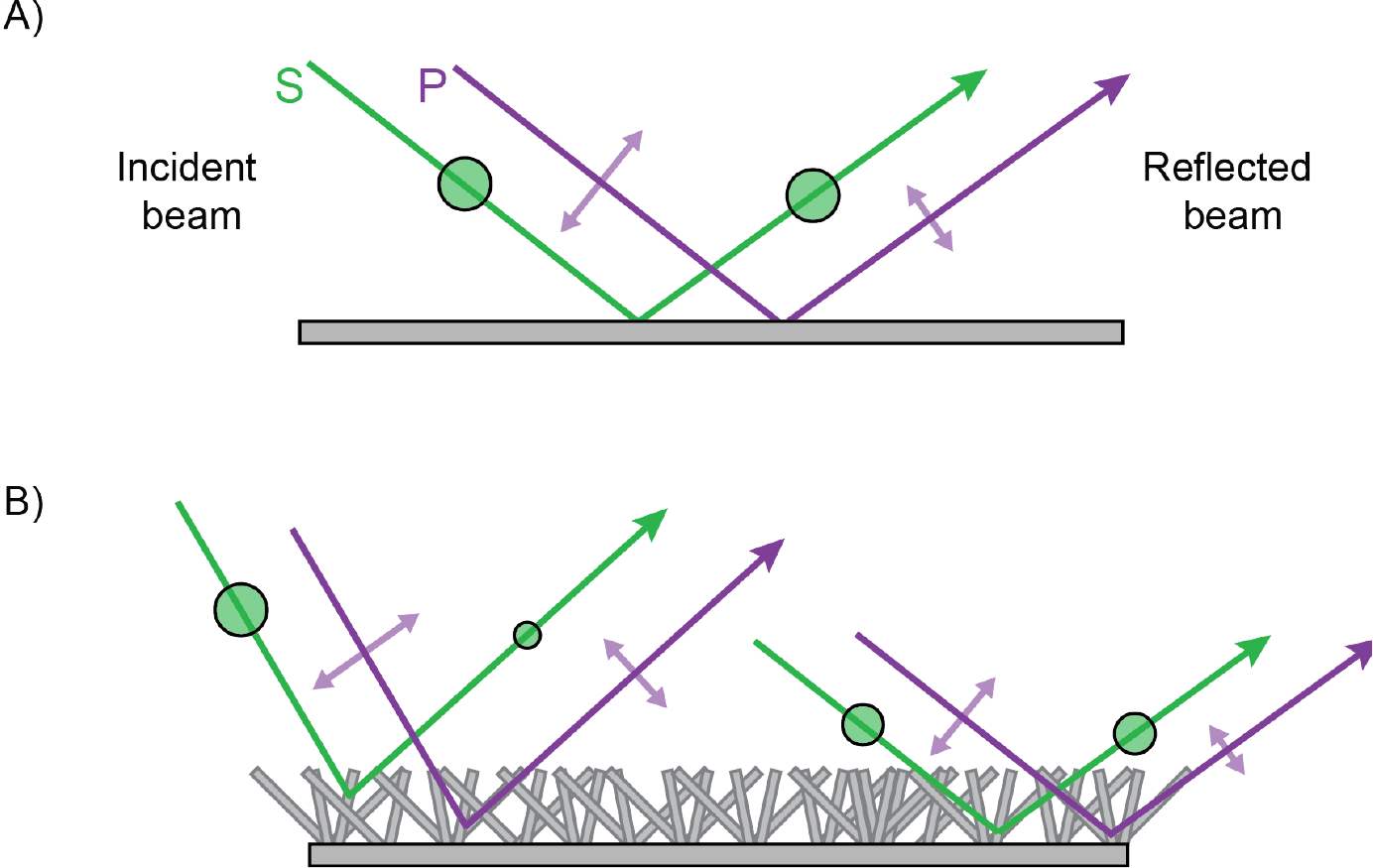
Comparison of reflected intensity of p-and s-polarized beams from a smooth (A) or fibrous material (B). (**A**) While a relatively large proportion of the intensity of s-polarized light (AoP into the page; intensity shown as size of the green circle) is preserved as the angle of reflection approaches Brewster’s angle (see §4.2.1), reflected intensity of p-polarized light (AoP in the plane of the page; intensity shown as arrow length) decreases as a function of increasing angle of incidence, reaching a minimum at Brewster’s angle. (**B**) The same effects also occur in ‘rough’ or fibrous surfaces, but an observer at a given angle to the surface sees light that has been reflected at a greater diversity of angles, including those angles for which intensity of reflected p-polarized light is high (left beam).

To further decrease the relative contribution of direct specular reflections from a surface, bright, diffuse ambient lighting can also be added to an experimental arrangement. The apparent lightness of the surface can be manipulated through the ratio of the ambient-and stimulus-light intensities, provided that the surfaces illuminated have high diffuse reflectance. We propose that, through careful consideration of the materials visible to an animal during experiments and calibration of sources of ambient lighting, confounding cues from surface reflections can be essentially eliminated under most circumstances.

#### 5.1.2 Viewing-Angle Effects in Polarizers

Laminated polarizers have been used to manipulate the AoP of stimulus light in most studies to date (see §4.1.1). As a stimulus, an unpolarized beam transmitted through a polarizer should be the same intensity regardless of that polarizer’s transmission axis (TA) orientation. This basic requirement is met by all commercially-available polarizers for a beam at normal incidence, but at non-normal incidence angles transmitted intensity varies as a function of TA orientation (Fig. 5.3). As a result of this ‘viewing-angle’ effect, even at relatively modest off-axis incidence angles, differences in transmitted intensity may be discriminable. For incidence angles ≥40° the Michelson contrast (a measure of pattern detectability) between two polarizers with perpendicular TAs could be above the detection threshold (Fig. 5.4) for a range of animal species (*e.g.* cat *Felis sylvestris:* Blake *et al.*, 1974; honeybee *Apis mellifera:* Bidwell & Goodman, 1993) and above 50° this difference would be detectable to most species capable of spatial vision (*e.g.* goldfish *Carassius auratus:* Northmore & Dvorak, 1979; various birds: Ghim & Hodos, 2006; Lind *et al.*, 2012). When two polarizers are presented to an animal simultaneously, the animal might therefore detect an intensity difference between them when viewing angle is not controlled.

**Fig. 5.3.**
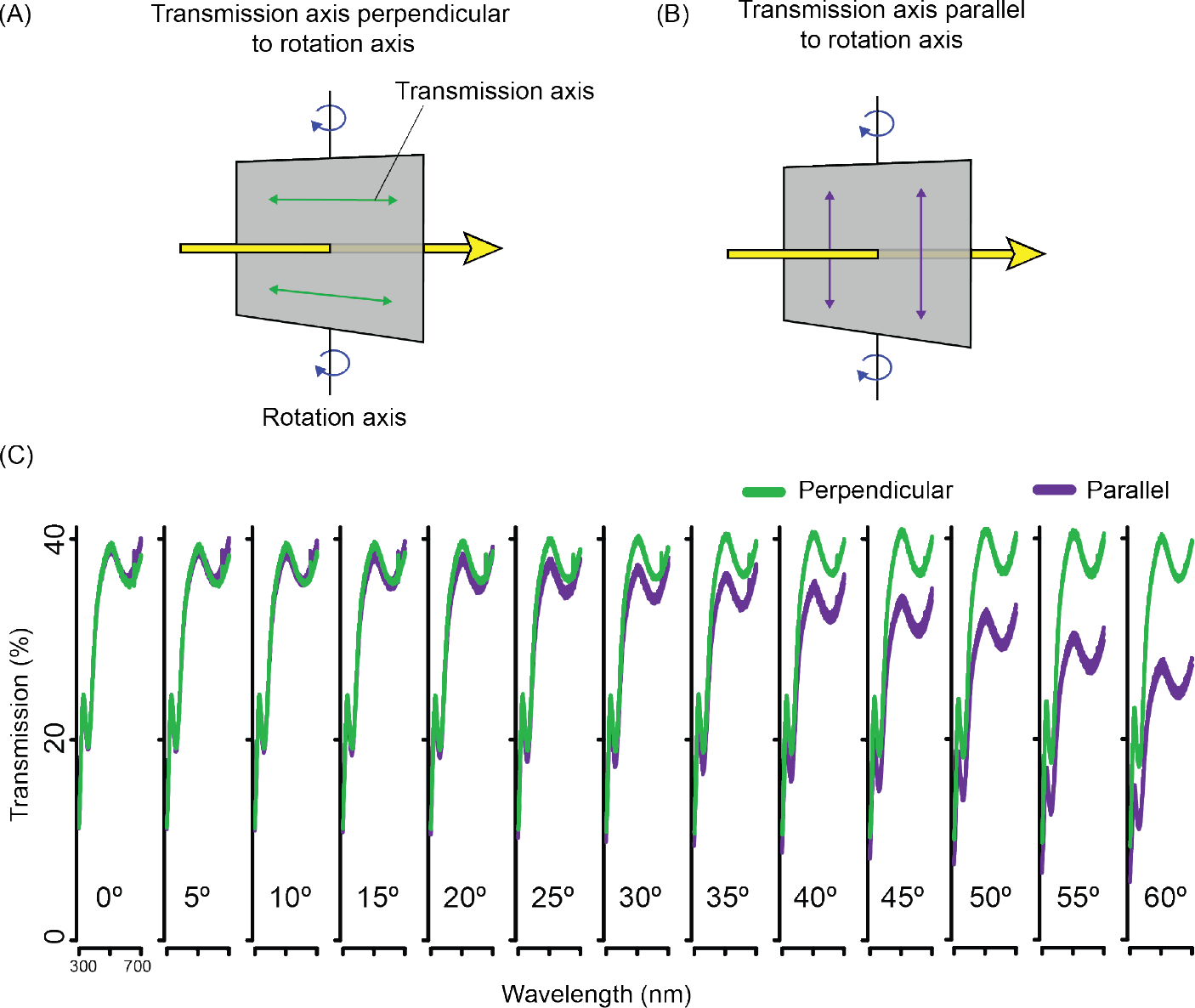
Changes in polarizer transmittance with angle of incidence. (**A**) The arrangement in which the polarizer’s transmission axis (TA) is perpendicular to the axis of rotation—relative to the incident beam (parallel to the plane of incidence). For this arrangement, transmission is not modulated as a function of angle of incidence. (**B**) The arrangement in which the polarizer’s TA is parallel to the axis of rotation. For this arrangement the proportion of the incident beam’s intensity that is transmitted decreases as a function of the angle of incidence (see panel C). (**C**) The transmittance spectrum of a UV-grade polarizer (HNP’B, Polaroid, USA) recorded at a range of incidence angles in the orientation described in (A-B). For incidence angles ≥ 25° there is a clear reduction in transmitted intensity when TA orientation is parallel to the axis of rotation (as compared with when the TA is perpendicular to this axis). For more details of the measurement apparatus see supplement S6.

**Fig. 5.4.**
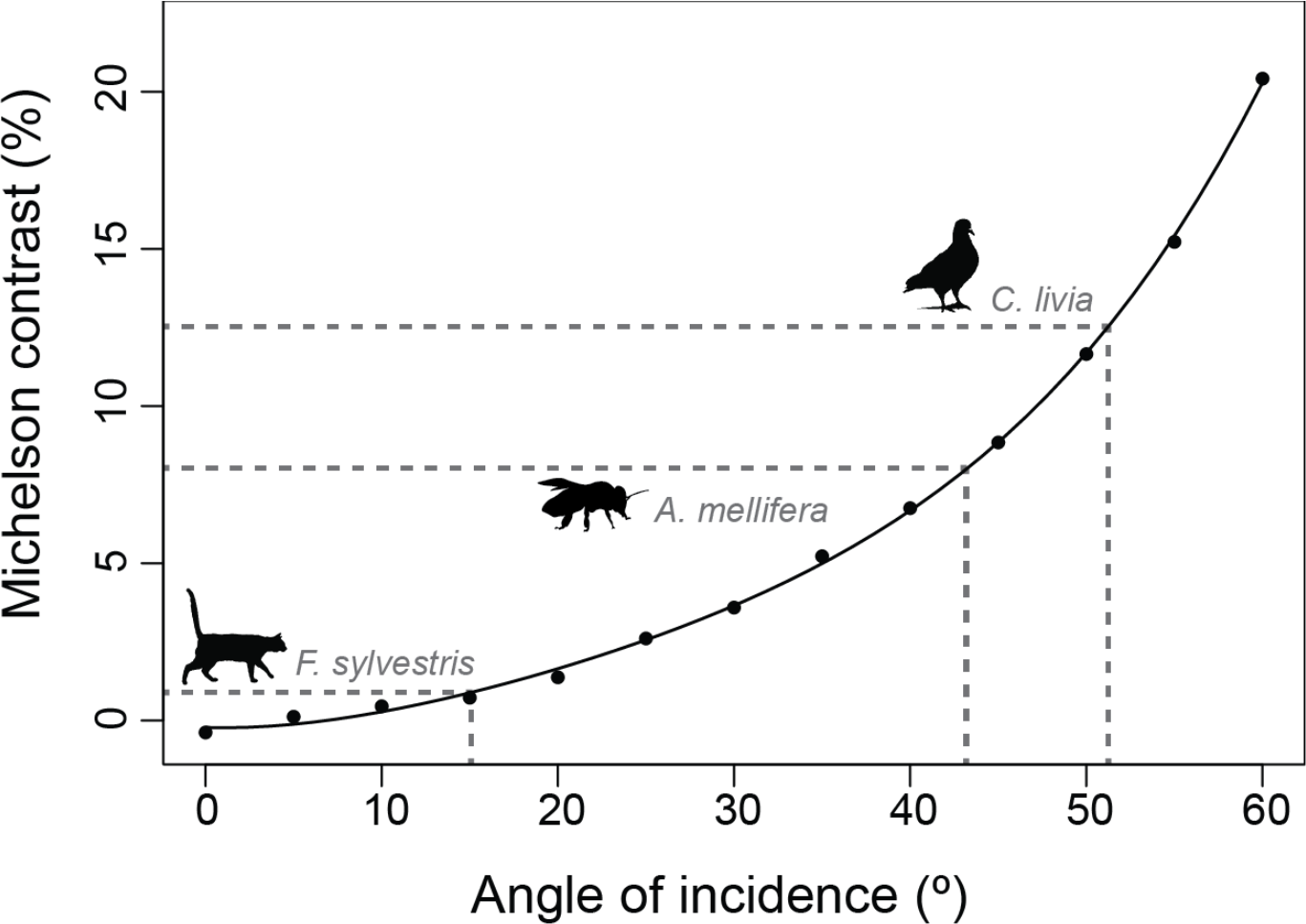
Intensity contrast between polarizer orientations. Michelson contrast in transmitted intensity between two adjacent polarizers with perpendicular transmission axes, at incidence angles 0°−60° (points: measured; line: fitted polynomial). Example maximum contrast sensitivity thresholds for model species (*Felis sylvestris*: Blake *et al.*, 1974; *Columba livia:* Ghim & Hodos, 2006; *Apis mellifera;* Bidwell & Goodman, 1993) provided for reference.

In addition to observing changes in transmitted intensity directly, light transmitted at large incidence angles may also illuminate surfaces within an experimental arrangement. Surfaces illuminated at large incidence angles may receive more or less illumination as a function of TA orientation. When displaying a polarized stimulus, in the form of a backlit polarizer, in a box-shaped experimental chamber (such as a Skinner box, y-maze or choice-chamber experiment), projected off-axis transmission can form a confounding brightness pattern (Fig. 5.5). As such, even when the animal’s viewing angle is restricted, intensity differences that result from transmission through a polarizer at large incidence angles may be present.

**Fig. 5.5.**
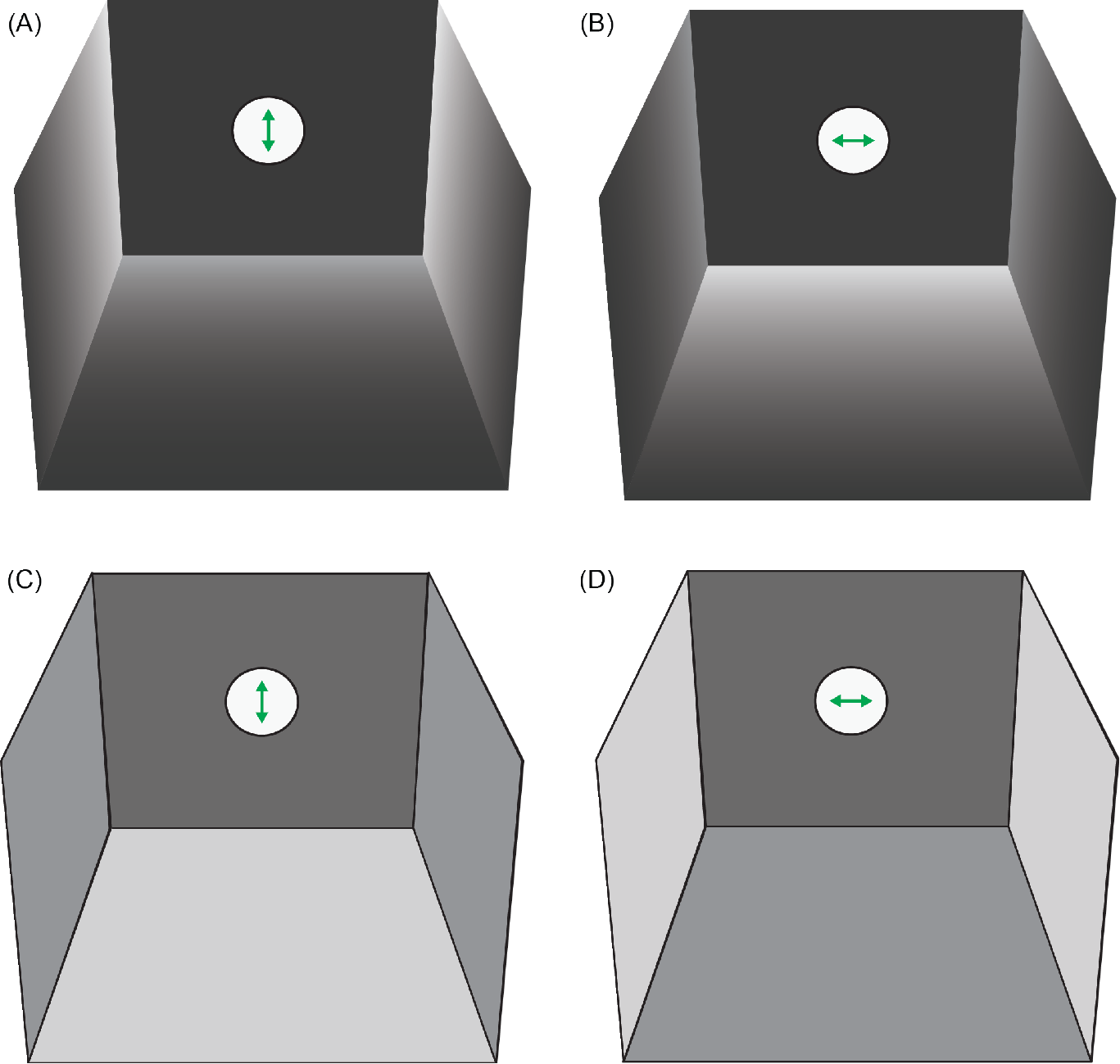
An illustrated example of a y-maze paradigm that might be confounded by polarized stimuli. In y-maze arm (**A**) vertically-polarized stimulus light reflected from the chamber walls is brighter than that reflected from the floor. In arm (**B**) stimulus light is horizontally-polarized and the situation is reversed, making average brightness of the arm as a whole lower than for (A). In (**C-D**) the arena is lined with a rougher substance with a high diffuse reflectance, minimising these differences. In this scenario, projected off-axis transmitted illumination may act as an intensity confound. In y-maze arm (**C**) the TA of the polarizer is vertical, and hence transmitted light that illuminates the chamber’s vertical walls is darker than that which illuminates the chamber’s horizontal floor. In y-maze arm (**D**) the polarizer’s TA is horizontally oriented, and the pattern is reversed.

### 5.2 Controlling for Intensity Confounds

#### 5.2.1 Restriction of Animal’s Viewing Angle

The effects discussed above can be addressed directly by restricting the angles at which the study animal views polarized stimuli and surfaces illuminated by them. In some cases it may be possible to measure the strength of surface-reflection cues across a range of angles, and the viewing angle effect for a given polarizer. These could be used to determine a range of viewing angles below the minimum discriminable contrast for the study species (see Fig. 5.4 for an example). This approach would, however, have limited applications for experimental paradigms requiring either broad-field stimuli or free movement of the study animal, and requires some prior estimate of contrast sensitivity.

#### 5.2.2 Collimated Stimulus Light

A simple means of limiting confounding intensity cues is to use a collimated light source. Collimated light may be produced via the introduction of a collimator, in the form of a lens or mirror, into the light path before the polarizer. The light beams then pass through the polarizer at close to normal incidence. This procedure has been used in a number of studies of polarization sensitivity (McCann & Arnett, 1972; Edrich & Helverson, 1976; Wolf *et al.*, 1980; Henze & Labhart, 2007), typically employing a light source-collimator-diffuser-polarizer light path (Fig. 5.6).

**Fig. 5.6.**
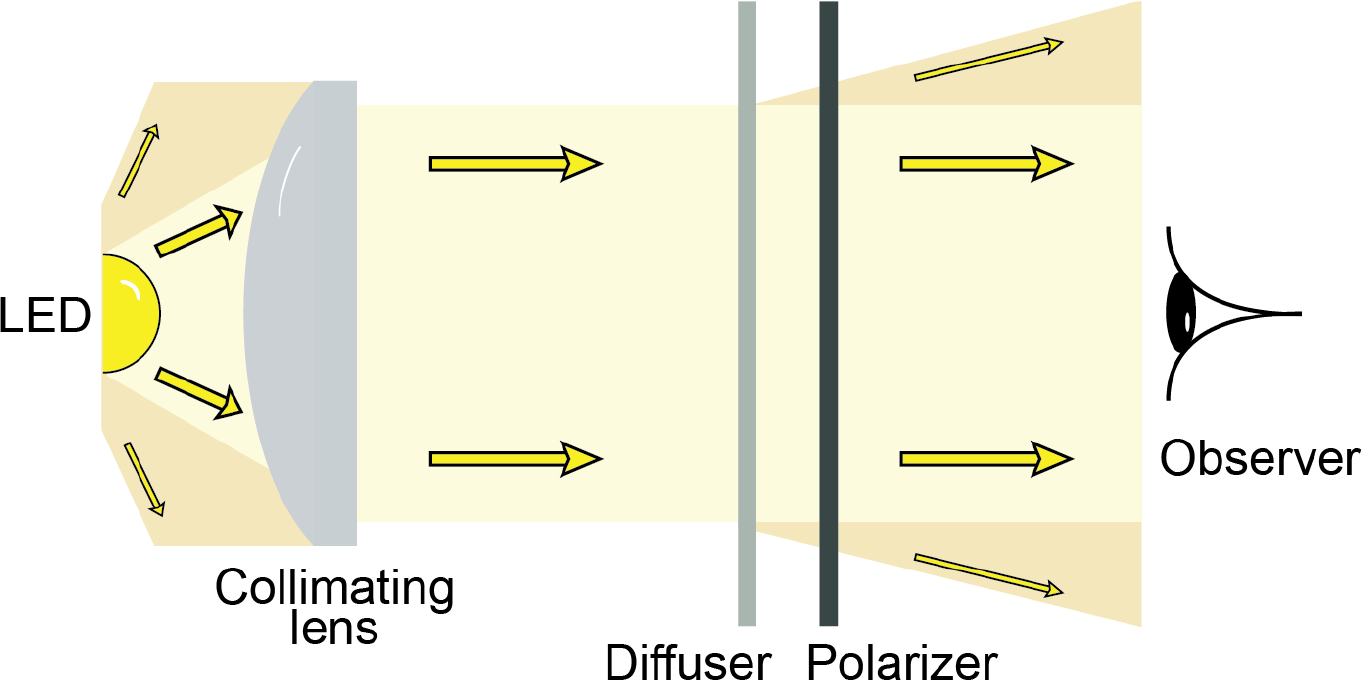
Collimating stimulus light. A collimating lens can be used to control the spread of a stimulus beam, so that the study animal observes stimulus light that is at normal incidence to the polarizer. Note that while some stimulus light is scattered by the diffuser, the spread is narrower than was the case for the initial light source.

This method does, however, introduce some restrictions on animal position and light source properties. Because collimated light enters and exits the polarizer at a narrow range of viewing angles, the stimulus is not visible to the study animal outside of this range. Therefore this technique is better suited to electrophysiological investigations (*e.g.* McCann & Arnett, 1972) or trackball behavioural paradigms (*e.g.* Henze & Labhart, 2007) in which the position of the animal is necessarily restricted throughout the experiment.

Restrictions in light source properties may also be problematic. Standard practice is to introduce a diffuser directly before the polarizer, to ensure that light incident on the polarizer is not already polarized, which would modulate its intensity. Diffusers scatter normally-incident light, reducing the effective collimation of light incident on the polarizer. In most cases, it may be practical to alter the order of optical components, so that the beam passes through the diffuser before the collimator.

## 6. Conclusions

While technology progresses and new techniques are developed, it continues to be challenging to control for confounding cues when presenting polarized stimuli to animals. We aim to give the reader an overview of the most common and effective techniques currently available, and to provide a toolkit for critically assessing the suitability of these experimental methods in a research setting; although the methods for the production and calibration of polarized stimuli presented in this review necessarily represents only a subset of the full variety of available techniques.

The polarization of light can be understood as several different properties of light, spatially and temporally averaged. Polarized light is abundant in nature, and it is the polarization states of these light sources that are the most biologically relevant. The terminology used to describe the qualities of a polarized light beam varies somewhat within the biological literature, and we recommend that researchers clearly specify a definition and how it is calculated, for example using terms such as angle of polarization (AoP) and degree of polarization (DoP).

Polarized stimuli should be measured to understand their appearance to the animal under study. This requires some form of light detector and polarizing filter, and can be achieved to different degrees of completeness across a given region in space and across the UV-visible spectrum; the most complete characterisation will often require more than one method. Measurement techniques should be chosen with reference to the aims of the experiment and what is known about the visual system of the the study species.

A range of methods are available for controlling the angle and degree of linear polarization of polarized stimuli. While the most commonly used methods, sheet polarizers and specularly reflecting surfaces, allow the production of stimuli that consist of a single AoP polarized to a high degree, recent advances make it possible to recreate more naturalistic and dynamic patterns of polarization under controlled conditions.

Confounding cues are the predominant challenge in planning and conducting animal experiments involving polarization. Their sources may be optical effects incidental to the design of the experimental apparatus, as well as insufficient control or calibration of the stimulus. We recommend a combination of control of the animal’s view of the stimulus, the stimulus’ properties themselves and careful measurement, as means to produce reliable, repeatable and meaningful results in polarization vision studies.

## Supplement

**S1.**
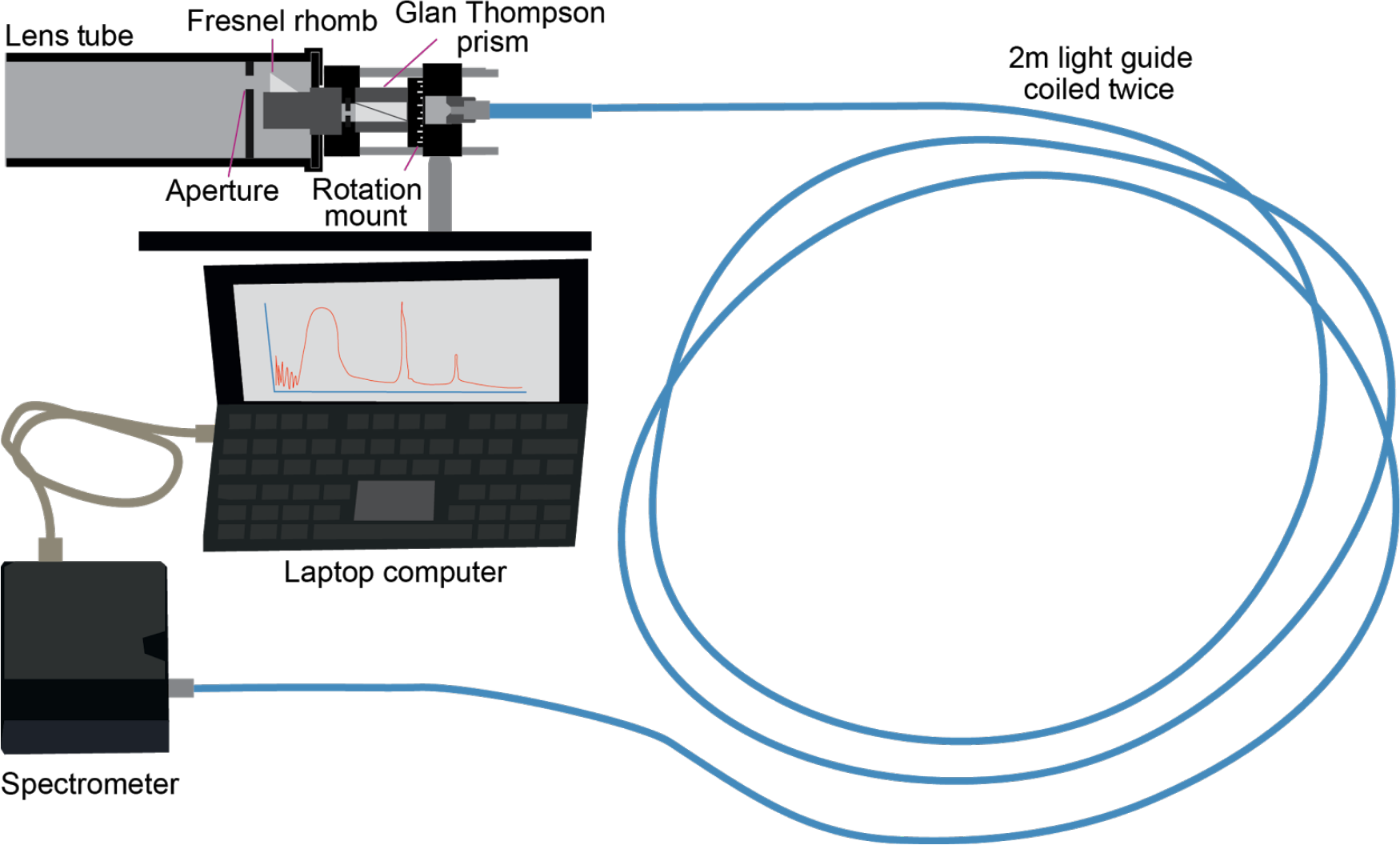
Example Arrangement for Spectral Characterisation of Polarization. In this arrangement the beam of light is analysed by polarizers placed in the light path before it enters the optical fibre (blue). An aperture and lens tube restrict the region in space sampled, as well as the range of off-axis incidence angles at which light enters the polarizers. The Fresnel rhomb, which converts circular polarization to linear polarization, displaces the light path, so this displacement needs to be compensated for by raising (or lowering) the whole apparatus between the measurements of the Stokes parameters’ linear (S1 and S2) and elliptical (S3) components. Alternatively, an additional component can be added to allow the whole apparatus to rotate around the light-collecting surface of the Fresnel rhomb (Gagnon & Marshall, 2016). In this case, when the Fresnel rhomb’s fast axis is aligned with the the Glan Thompson’s transmission axis, the linear component is analysed, and when the Glan Thompson prism’s transmission axis is at ±45° the elliptical component is measured. Light relayed to the spectrometer by a coiled light guide should be depolarized, so no introduced polarization affects measurements (see S2). Since only the ratio of the intensities recorded for each polarizer orientation (see §2.1.2) is required to calculate the polarization state, rather than their precise radiometric values, the spectrometer does not need to be calibrated for spectral radiance measurements. Each measurement can be recorded as the number of spectrometer counts at each wavelength for a given integration time.

**S2.**
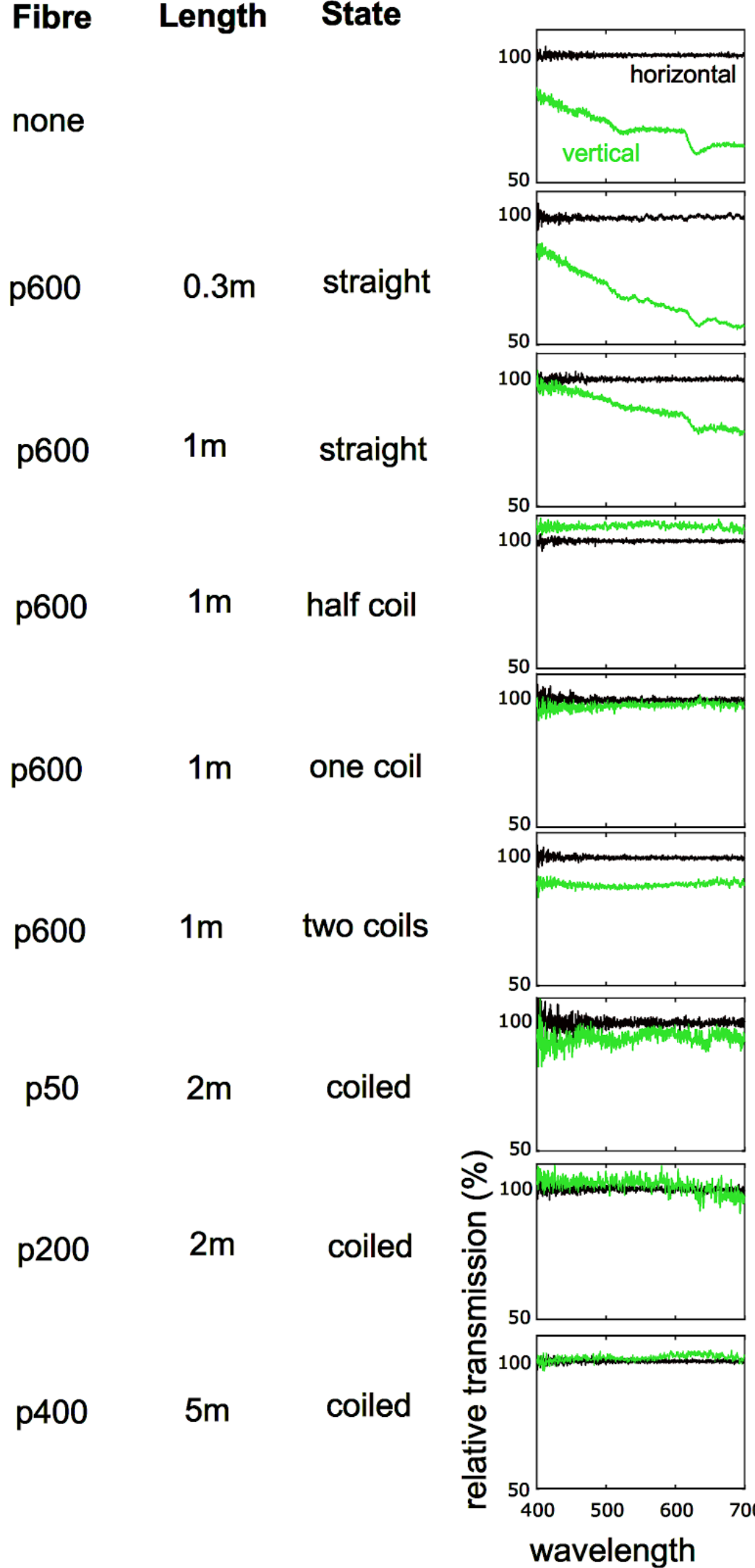
Polarization Sensitivity in a Spectrometer. Due to optical effects such as mode propagation, it is possible that light with certain polarization characteristics (*i.e.* specific angles) is propagated with less loss than other polarization states. To check for this effect, record the intensity of light passing through a rotating linear polarizer in front of an unpolarized light source with a bare fibre. The spectrum recorded by the device should remain constant regardless of the angle of incoming polarization. Where polarization sensitivity has not been eliminated, periodic fluctuations appear in the spectral measurement. This inherent polarization sensitivity can be overcome by ensuring that a long (>1m) fibre is used and also by coiling the fibre multiple times. *N.B.* optic fibres are supplied with a recommended minimum bend radius, which should be avoided.

**S3.**
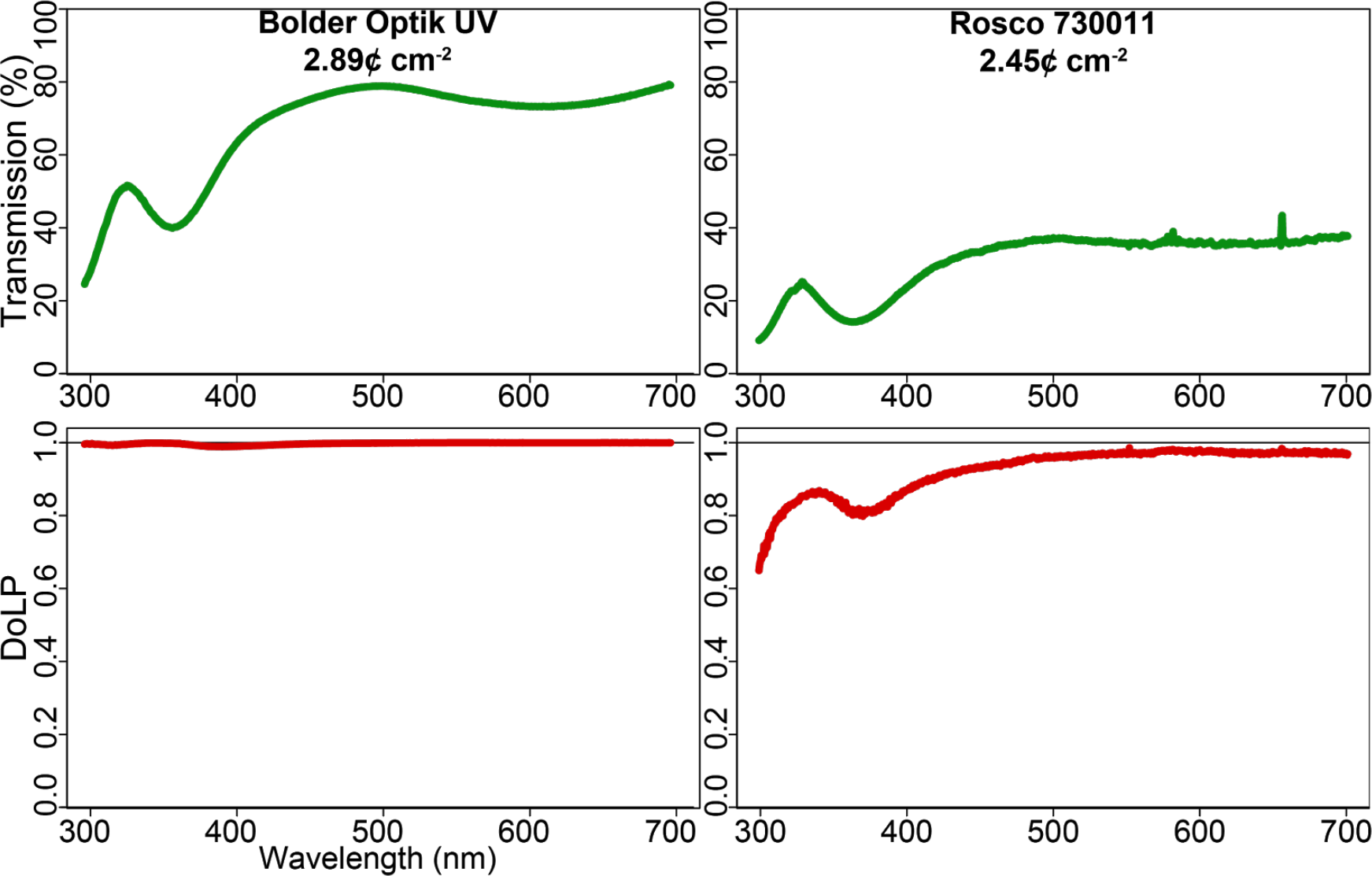
Example Polarizer Transmission Spectra and Prices. Transmission (**top**) and degree of linear polarization (**bottom**) spectra for a technical grade (**left**: Bolder Vision Optik, USA) and theatrical grade (**right**: RoscoLab Ltd., UK) polarizing sheet. Prices in USD/100 per square centimeter based on unit prices for intact sheets (Bolder: 1000 × 620 mm; Rosco: 510 × 430 mm) excluding sales taxes and shipping costs.

**S4.**
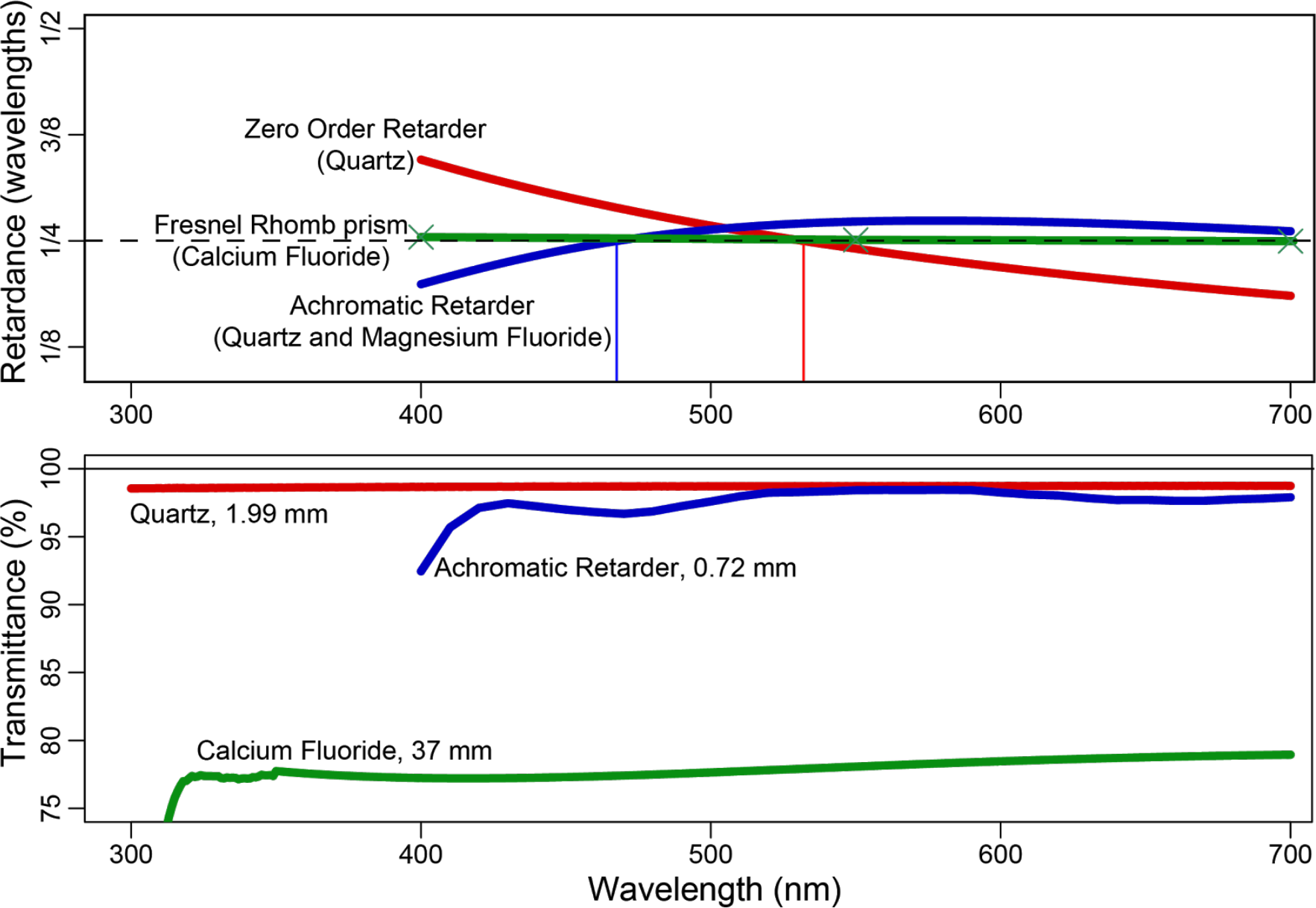
Retardation Devices—Retardance and Transmittance. Retardance **(top)** and transmission **(bottom)** spectra for three types of optical-grade quarter wave retarder. Zero order retarders (red line; WPQ05M-532: Thorlabs GmbH, Germany) slow one component of linear polarization by a given distance (across most of the spectrum) and therefore only retard by one quarter of a specific wavelength, as stated by the manufacturer (here 532 nm). ‘Achromatic’ retarders
(blue line; AQWP05M-600: Thorlabs) retard the component parallel to their slow axis by approximately one quarter of a wavelength across a wider range. One advantage of zero-order and achromatic retarders is that they can be relatively thin, preserving the spectrum of transmitted light (**bottom**). Fresnel rhombs (green; FR600QM: Thorlabs; crosses: measured; line: fitted curve) retard by almost exactly one quarter of a wavelength across their transmission spectrum, but the prism’s shape requires the light path to be longer than for a zero-order or achromatic retarder with an equivalent collection area. All retardance measurements and material transmission spectra from www.thorlabs.com, transmittance spectra of individual retarders estimated from material transmission spectra and light path distance through the retarder for WPQ05M-532 and FR600QM.

**S5.**
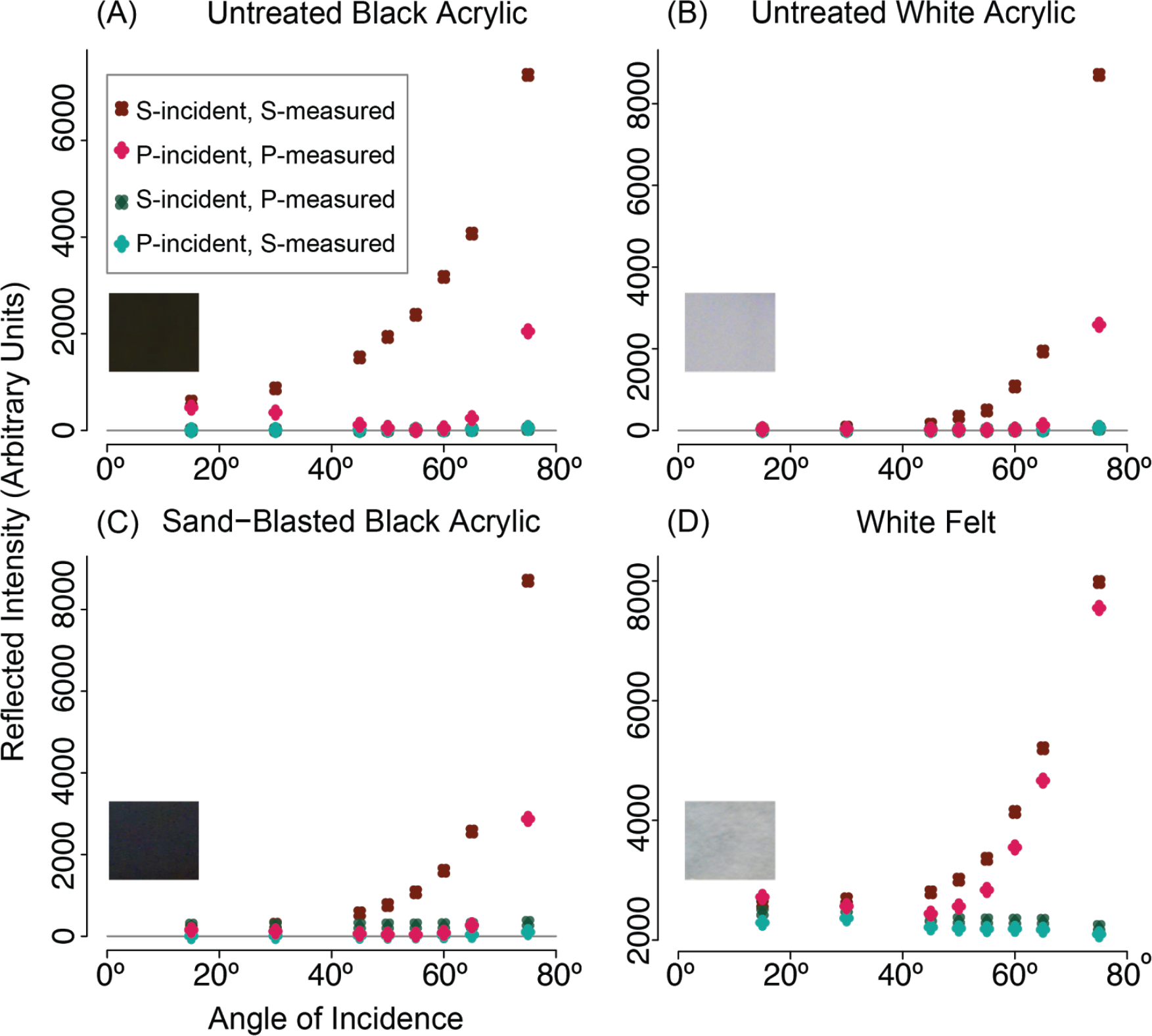
Measurements of Reflected s-and p-Polarization. (**A**) Black acrylic (Perspex, Weybridge, UK). For angles of incidence 45-60° the intensity of p-polarized reflected light is near zero, as a proportion of the reflected intensity of s-polarized light. *N.B.* When incident light is s-polarized, none of its reflected intensity is p-polarized, and when incident light is p-polarized reflected light is not s-polarized. (**B**) White acrylic (Perspex, UK). The ratio of reflected s-polarized light to p-polarized light is lower than for black acrylic. (**C**) The same block of black acrylic as (A.), sand-blasted to create a ‘rough’ surface. The increase in ratio with angle of incidence is lower than for untreated black acrylic. (**D**) White felt (Fabric Land Ltd., UK). This material is both highly reflective and ‘rough’ (fibrous), and hence the ratio of reflected s-polarization to p-polarization is low at all angles. Retardation by thin fibres may be responsible for the apparent conversion of s-polarized light to p-polarized, and p-polarized to s-polarized (green squares and rhombuses respectively). *N.B.* Reflected intensity falls within a narrower range in (D) than in the preceding panels.

**S6.**
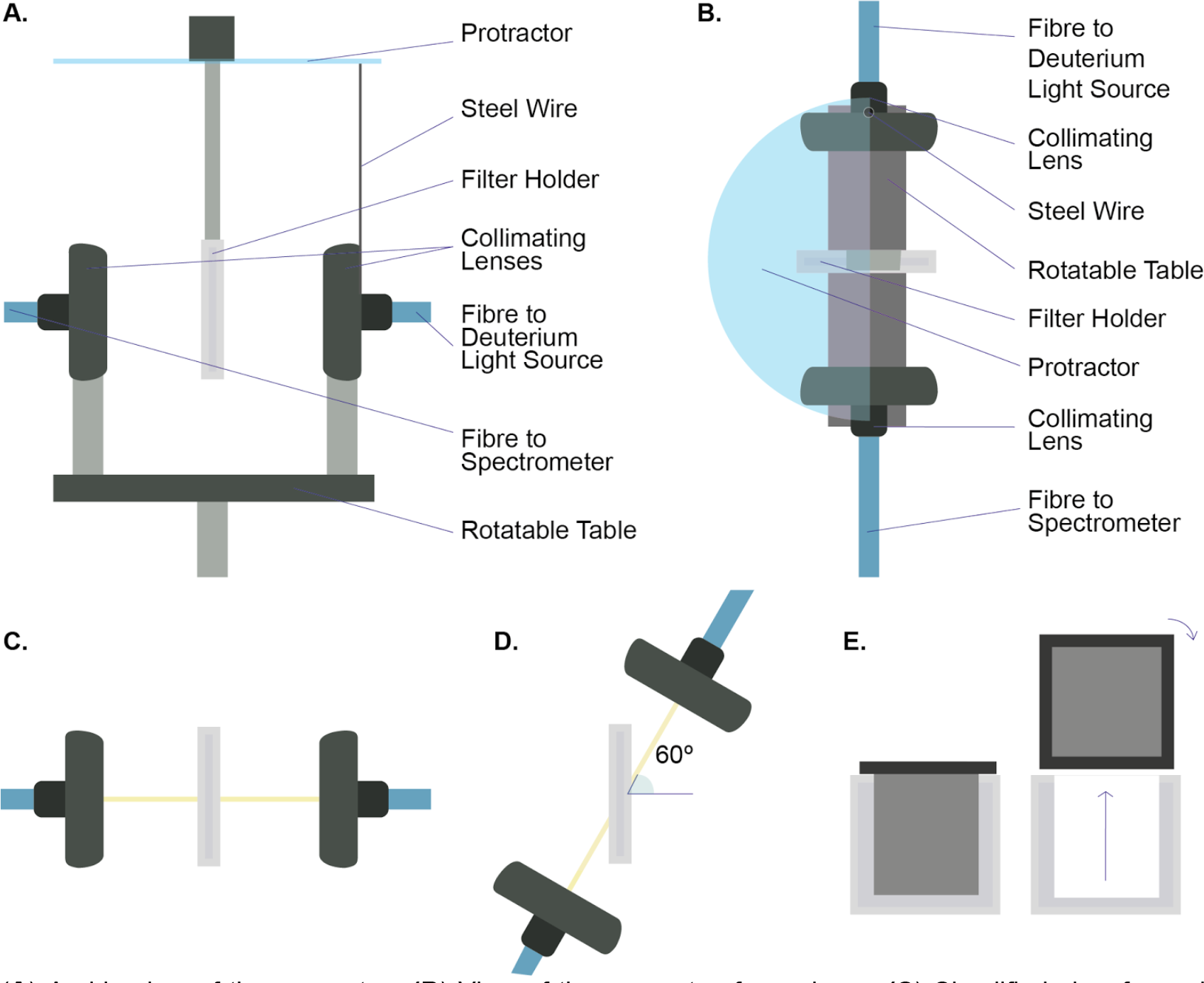
Goniometric Apparatus for Measuring Polarizer Transmission. (**A**) A side view of the apparatus. (**B**) View of the apparatus from above. (**C**) Simplified view from above showing the light path (yellow) at an incidence angle of 0°. (**D**) The arrangement and light path for an incidence angle of 60°. (**E**) Front view of the filter holder, from which the polarizer was removed and rotated by 90° between measurements. In order to compare differences in transmission through two perpendicular TA orientations as a function of incidence angle, an apparatus was developed that allowed the projection of light through a polarizer at a range of incidence angles. Broad spectrum light from a calibrated light source (DT-MINI-2-GS; Ocean Optics Inc., FL, USA) was projected through a collimating lens (74-UV; Ocean Optics Inc.), focussed to infinity, onto the polarizer. Transmitted light was collected by another collimating lens, equidistant from the polarizer, and relayed to a spectrometer (QE65000; Ocean Optics). Five sets of measurements across each incidence angle were recorded with the TA orientated perpendicular to the rotation axis (horizontal TA as viewed in (E)), with the TA oriented parallel to the rotation axis (vertical TA) and with no polarizer (in order to calculate transmittance).

**S7.**
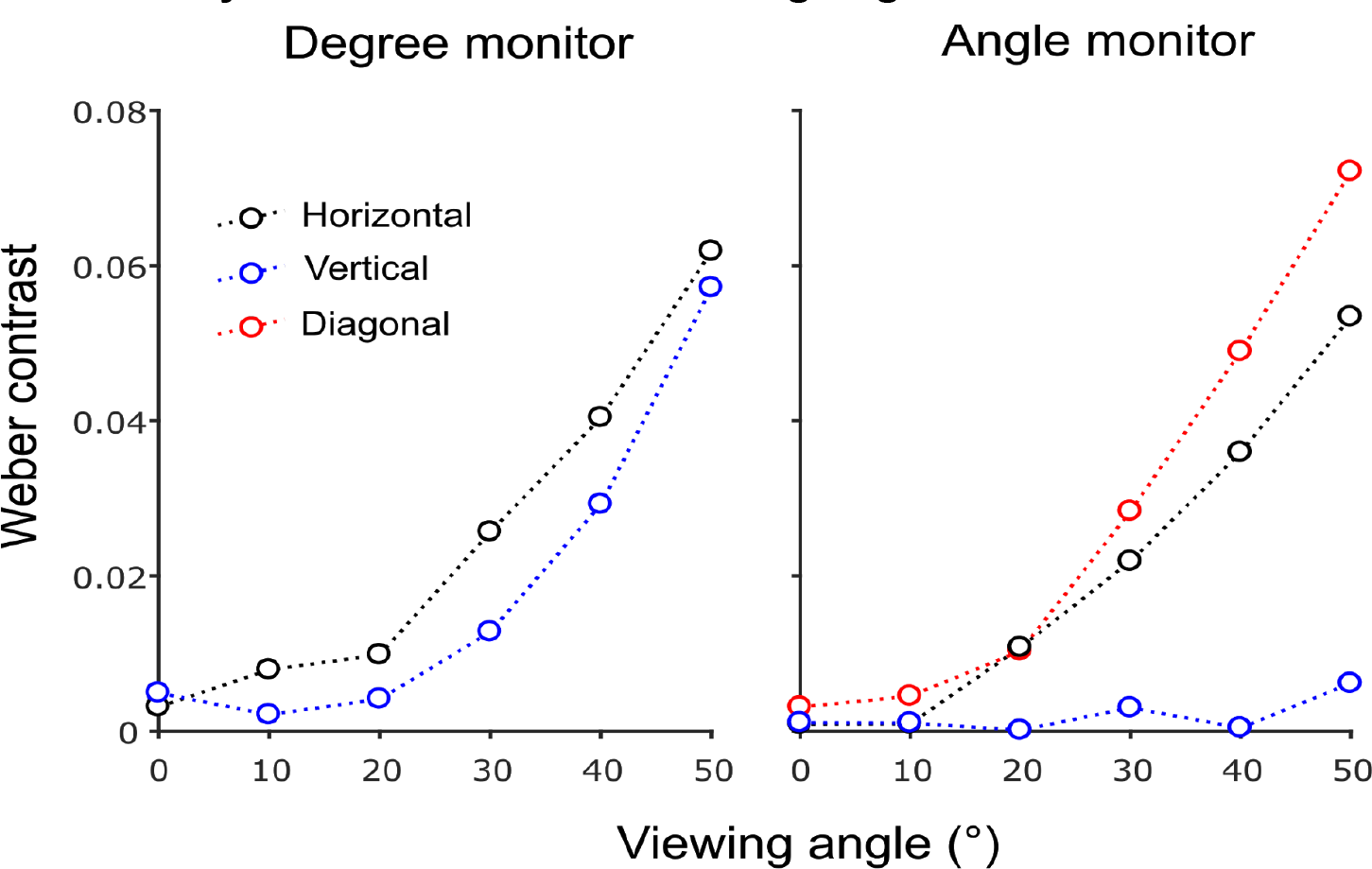
Intensity artifacts at off-axis viewing angles in LCD monitors. Contrast between the maximum and minimum voltage states (RGB values of 0 or 255; white or black in an unmodified LCD monitor) at off-axis viewing angles in the horizontal (black points) and vertical (blue points) planes, for an LCD monitor modified to control degree of polarization (**left;** 1905FP, Dell) and one modified to control angle of polarization (**right**; 1908fpc, Dell). Each measurement was taken through a coiled 2m, 400μm diameter fibre with a s≤3° gershun tube, connected to a spectrometer (Flame, Ocean Optics). Since for the “angle” monitor the two states have ≈0 contrast for vertical off-axis viewing angles, a further set of measurements were made in the plane diagonal to the screen (red points).

**S8.**
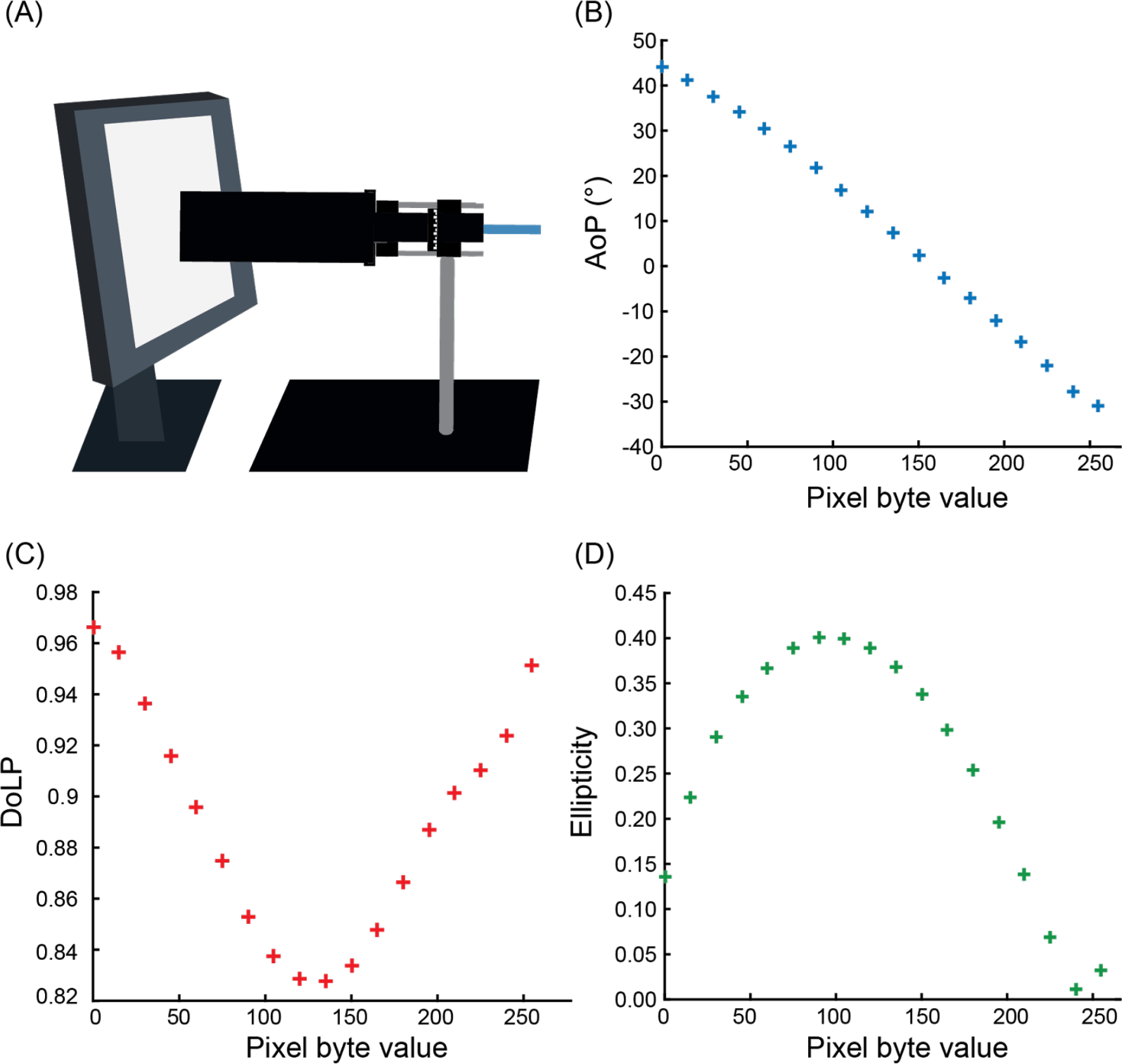
Calibration of LCD Monitors. As described in section 4, an LCD monitor may be converted to a polarization monitor by removing the frontmost polarizer, after which the polarization of the monitor’s display may be controlled via a computer using the 256-level (8-bit) RGB scale. In order to understand how the specific brand and model of monitor alters the polarization of transmitted light for different RGB values, it is absolutely essential to measure the angle and degree of polarization as well as the ellipticity of light from the monitor at a particular RGB value using one of the methods described in section 3 (Fig. 5.7A). The optical properties of the monitor can vary considerably as a function of wavelength and RGB value, so it may be necessary to measure polarization across the spectrum (see §3.2.2) and to sample at frequent intervals across the 8-bit range. Where wavelength dependence becomes problematic, it may be avoided by replacing the monitor’s backlight with a custom light source, such as a bank of LEDs with a narrow emission spectrum (see §4.1.3).Once steps have been taken to avoid RGB values between which polarization varies excessively, an experimenter may present an animal with a complex dynamic stimulus composed of regions with different polarization characteristics with little to no intensity variation. Measurement and calibration procedure for an LCD monitor. (**A**) Schematic showing arrangement of polarimetry equipment. (**B-D**) Examples of measurements for each pixel byte (RGB) value. The change in angle and the degree of polarization across the 256-level greyscale on a modified TN-film (AoP) monitor, (VL-15EC6, Viglen ltd., UK). This monitor can produce a ≈80° range of angles of polarization. At pixel byte values 0 and 210, angle of polarization differs by ≈78°, while remaining close in degree of linear polarization (**C**), so these two values could be chosen to create a pattern contrasting in angle of polarization.

## References

Able, K. P., & Able, M. A. (1993). Daytime calibration of magnetic orientation in a migratory bird requires a view of skylight polarization. Nature, 364(August), 523–525

Bainbridge, R., & Waterman, T. H. (1957). Polarized light and the orientation of two marine Crustacea. J. Exp. Biol., 34, 342–364.

Bidwell, N. J., & Goodman, L. J. (1993). Possible Functions of a Population of Descending Neurons in the Honeybees Visuo-Motor Pathway. Apidologie, 24(3), 333–354. doi:10.1051/apido:19930311

Blake, R., Cool, S. J., & Crawford, M. L. J. (1974). Visual resolution in the cat. Vision Res., 14(11), 1211–1217. doi:10.1016/0042-6989(74)90218-1

Boal, J. G., Shashar, N., Grable, M. M., Vaughan, K. H., Loew, E. R., & Hanlon, R. T. (2004). Behavioral Evidence for Intraspecific Signaling with Achromatic and Polarized Light by Cuttlefish (Mollusca: Cephalopoda). Behaviour, 141(7), 837–861.

Boda, P., Horváth, G., Kriska, G., Blahó, M., & Csabai, Z. (2014). Phototaxis and polarotaxis hand in hand: night dispersal flight of aquatic insects distracted synergistically by light intensity and reflection polarization. Naturwissenschaften, 101(5), 385–395. doi:10.1007/s00114-014-1166-2

Brewster, D. (1815). On the Laws Which Regulate the Polarisation of Light by Reflexion from Transparent Bodies. Phil. Trans. R. Soc., 105, 125–159.

Calabrese, G. M., Brady, P. C., Gruev, V., & Cummings, M. E. (2014). Polarization signaling in swordtails alters female mate preference. PNAS, 111(37), 13397–13402. doi:10.1073/pnas.1321368111

Cartron, L., Josef, N., Lerner, A., McCusker, S. D., Darmaillacq, A.-S., Dickel, L., & Shashar, N. (2013). Polarization vision can improve object detection in turbid waters by cuttlefish. J. Exp. Mar. Biol. Ecol., 447, 80–85. doi:10.1016/j.jembe.2013.02.013

Chiou, T.-H., Kleinlogel, S., Cronin, T. W., Caldwell, R., Loeffler, B., Siddiqi, A., … Marshall, N. J. (2008). Circular polarization vision in a stomatopod crustacean. Curr. Biol., 18(6), 1–6. doi:10.1016/j.cub.2008.02.066

Chiou, T.-H., Marshall, N. J., Caldwell, R. L., & Cronin, T. W. (2011). Changes in light-reflecting properties of signalling appendages alter mate choice behaviour in a stomatopod crustacean Haptosquilla trispinosa. Mar. Freshw. Behav. Physiol., 44(1), 1–11. doi:10.1080/10236244.2010.546064

Chiou, T.-H., Mathger, L. M., Hanlon, R. T., & Cronin, T. W. (2007). Spectral and spatial properties of polarized light reflections from the arms of squid (Loligo pealeii) and cuttlefish (Sepia officinalis L.). J. Exp. Biol., 210, 3624–3635. doi:10.1242/jeb.006932

Chiou, T.-H., Place, A. R., Caldwell, R. L., Marshall, N. J., & Cronin, T. W. (2012). A novel function for a carotenoid: astaxanthin used as a polarizer for visual signalling in a mantis shrimp. J. Exp. Biol., 215, 584–589. doi:10.1242/jeb.066019

Coemans, M. A., Vos Hzn, J. J., & Nuboer, J. F. (1990). No evidence for polarization sensitivity in the pigeon. Naturwissenschaften, 77(3), 138–142.

Coffin, D. (1997). DCRAW: Decoding Raw Digital Photos. http://www.cybercom.net/~dcoffin/dcraw/.

Cronin, T. W., & Shashar, N. (2001). The linearly polarized light field in clear, tropical marine waters: spatial and temporal variation of light intensity, degree of polarization and e-vector angle. J. Exp. Biol., 204, 2461–2467.

Cronin, T. W., Shashar, N., Caldwell, R. L., Marshall, N. J., Cheroske, A. G., & Chiou, T.-H. (2003). Polarization Vision and Its Role in Biological Signaling. Integ. Comp. Biol., 43(4), 549–558. l

Cronin, T. W., Warrant, E. J., & Greiner, B. (2006). Celestial polarization patterns during twilight. Appl. Opt., 45(22), 5582–5589. doi:10.1364/A0.45.005582

Cronin, T. W., Chiou, T., Caldwell, R. L.-H., Roberts, N. W., & Marshall, N. J. (2009). Polarization signals in mantis shrimps. In J. A. Shaw & J. S. Tyo (Eds.), Proceedings of SPIE (p. 74610C). doi:10.1117/12.828492

Csabai, Z., Boda, P., Bernȧth, B., Kriska, G., & Horváth, G. (2006). A ‘polarisation sun-dial’ dictates the optimal time of day for dispersal by flying aquatic insects. Freshw. Biol., 51(7), 1341–1350. doi:10.1111/j.1365-2427.2006.01576.x

Dacke, M., Byrne, M. J., Scholtz, C. H., & Warrant, E. J. (2004). Lunar orientation in a beetle. Proc. R. Soc. B, 271, 361–365. doi:10.1098/rspb.2003.2594

Daly, I. M., How, M. J., Partridge, J. C., Temple, S. E., Marshall, N. J., Cronin, T. W., & Roberts, N. W. (2016). Dynamic polarization vision in mantis shrimps. Nat. Commun., 7(12140). doi:10.1038/ncomms12140

Edrich, W., & Helverson, O. von. (1976). Polarized light orientation of the honey bee: the minimum visual angle. J. Comp. Physiol. A, 109(3), 309–314. doi:10.1007/BF00663611

Egri, A., Blahó, M., Kriska, G., Farkas, R., Gyurkovszky, M., Akesson, S., & Horváth, G. (2012). Polarotactic tabanids find striped patterns with brightness and/or polarization modulation least attractive: an advantage of zebra stripes. J. Exp. Biol., 215, 736–745. doi:10.1242/jeb.065540

Egri, Á., Farkas, A., Kriska, G., & Horváth, G. (2016). Polarization sensitivity in Collembola: an experimental study of polarotaxis in the water-surface-inhabiting springtail, Podura aquatica. J. Exp. Biol., 219, 2567–2576. doi:10.1242/jeb.139295

el Jundi, B., Warrant, E. J., Byrne, M. J., Khaldy, L., Baird, E., Smolka, J., & Dacke, M. (2015). Neural coding underlying the cue preference for celestial orientation. PNAS, 112(36), 11395–11400. doi:10.1073/pnas.1501272112

Feller, K. D., Jordan, T. M., Wilby, D., & Roberts, N. W. (2017). Selection of the intrinsic polarization properties of animal optical materials creates enhanced structural reflectivity and camouflage. Phil. Trans. R. Soc. B, 372(1724), 2016–0336. doi:10.1098/rstb.2016.0336

Foster, J. J., Sharkey, C. R., Gaworska, A. V. A., Roberts, N. W., Whitney, H. M., & Partridge, J. C. (2014). Bumblebees Learn Polarization Patterns. Curr. Biol., 24(12), 1415–1420. doi:10.1016/j.cub.2014.05.007

Frisch, K. von. (1949). Die Polarisation des Himmelslichtes als orientierender Faktor bei den Tänzen der Bienen. Experientia, V/4, 142–8.

Gagnon, Y. L., & Marshall, N. J. (2016). Intuitive representation of photopolarimetric data using the polarization ellipse. J. Exp. Biol., 219, 2430–2434. doi:10.1242/jeb.139139

Gagnon, Y. L., Templin, R. M., How, M. J., & Marshall, N. J. (2015). Circularly polarized light as a communication signal in mantis shrimps. Curr. Biol., 25(23), 3074–3078. doi:10.1016/j.cub.2015.10.047

Gál, J., Horváth, G., Barta, A., & Wehner, R. (2001). Polarization of the moonlit clear night sky measured by full□sky imaging polarimetry at full Moon: Comparison of the polarization of moonlit and sunlit skies. J. Geophys. Res., 106(D19), 22647–22653. doi:10.1029/2000JD000085

Ghim, M. M., & Hodos, W. (2006). Spatial contrast sensitivity of birds. J. Comp. Physiol. A, 192(5), 523–534. doi:10.1007/s00359-005-0090-5

Glantz, R. M., & Schroeter, J. P. (2006). Polarization contrast and motion detection. J. Comp. Physiol. A, 192(9), 905–914. doi:10.1007/s00359-006-0127-4

Goddard, S. M., & Forward, R. B. (1991). The role of the underwater polarized light pattern, in sun compass navigation of the grass shrimp, Palaemonetes vulgaris. J. Comp. Physiol. A, 169(4), 479–491. doi:10.1007/BF00197660

Haidinger, W. von. (1844). Ueber das directe Erkennen des polarisirten Lichts und der Lage der Polarisationsebene. Annalen Der Physik, LXIII(9), 29–39.

Hawryshyn, C. W., & Bolger, A. E. (1990). Spatial orientation of trout to partially polarized light. J. Comp. Physiol. A, 167(5), 691–697.

Henze, M. J., & Labhart, T. (2007). Haze, clouds and limited sky visibility: polarotactic orientation of crickets under difficult stimulus conditions. J. Exp. Biol., 210, 3266–3276. doi:10.1242/jeb.007831

Horváth, G., Barta, A., & Hegedüs, R. (2014). Polarization of the Sky. In Polarized Light in Animal Vision (G. Horváth, Ed.). Berlin/Heidelberg: Springer. doi:10.1007/978-3-642-54718-8

Horváth, G., & Kriska, G. (2008). Polarization Vision in Aquatic Insects and Ecological Traps for Polarotactic Insects. In Aquatic insects: challenges to populations: proceedings of the Royal Entomological Society’s 24th symposium (J. Lancaster & R. A. Briers: Eds.), (pp. 204–229). CABI.

Horváth, G., Malik, P., Kriska, G., & Wildermuth, H. (2007). Ecological traps for dragonflies in a cemetery: the attraction of Sympetrum species (Odonata: Libellulidae) by horizontally polarizing black gravestones. Freshw. Biol., 52(9), 1700–1709. doi:10.1111/j.1365-2427.2007.01798.x

Horváth, G., & Varjú, D. (2004). Polarized Light in Animal Vision. Heidelberg/New York: Springer-Verlag.

Horváth, G., & Zeil, J. (1996). Kuwait oil lakes as insect traps. Nature, 379(January), 303–305.

How, M. J., Christy, J. H., Temple, S. E., Hemmi, J. M., Marshall, N. J., & Roberts, N. W. (2015). Target detection is enhanced by polarization vision in a fiddler crab. Curr. Biol., 25(23), 3069–3073. doi:10.1016/j.cub.2015.09.073

How, M. J., Christy, J., Roberts, N. W., & Marshall, N. J. (2014). Null point of discrimination in crustacean polarisation vision. J. Exp. Biol., 217, 2462–7. doi:10.1242/jeb.103457

How, M. J., Porter, M. L., Radford, A. N., Feller, K. D., Temple, S. E., Caldwell, R. L., … Roberts, N. W. (2014). Out of the blue: The evolution of horizontally polarized signals in Haptosquilla (Crustacea, Stomatopoda, Protosquillidae). J. Exp. Biol., 2017, 3425–3431. doi:10.1242/jeb.107581

How, M. J., & Marshall, N. J. (2014). Polarization distance: a framework for modelling object detection by polarization vision systems. Proc. R. Soc. B, 281(2013–1632).

How, M. J., Pignatelli, V., Temple, S. E., Marshall, N. J., & Hemmi, J. M. (2012). High e-vector acuity in the polarisation vision system of the fiddler crab Uca vomeris. J. Exp. Biol., 215(Pt 12), 2128–34. doi:10.1242/jeb.068544

Jander, R., Daumer, K., & Waterman, T. H. (1963). Polarized light orientation by two Hawaiian decapod cephalopods. J. Comp. Physiol. A, 46(4), 383–394.

Johnsen, S. (2012). The Optics of Life: A Biologist’s Guide to Light in Nature. Princeton University Press.

Johnsen, S., Marshall, N. J., & Widder, E. A. (2011). Polarization sensitivity as a contrast enhancer in pelagic predators: lessons from in situ polarization imaging of transparent zooplankton. Phil. Trans. R. Soc. B, 366(1565), 655–670. doi:10.1098/rstb.2010.0193

Jordan, T. M., Partridge, J. C., & Roberts, N. W. (2012). Non-polarizing broadband multilayer reflectors in fish. Nat. Photonics, 6(11), 759–763. doi:10.1038/nphoton.2012.260

Jordan, T. M., Partridge, J. C., & Roberts, N. W. (2014). Disordered animal multilayer reflectors and the localization of light. J. R. Soc. Interface, 11, 2014–0948. doi:10.1098/rsif.2014.0948

Jordan, T. M., Wilby, D., Chiou, T.-H., Feller, K. D., Caldwell, R. L., Cronin, T. W., & Roberts, N. W. (2016). A shape-anisotropic reflective polarizer in a stomatopod crustacean. Sci. Rep., 6(21744). doi:10.1038/srep21744

Kelber, A. (1999). Why “false” colours are seen by butterflies. Nature, 402(6759), 251. doi:10.1038/46204

Kinoshita, M., Yamazato, K., & Arikawa, K. (2011). Polarization-based brightness discrimination in the foraging butterfly, Papilio xuthus. Phil. Trans. R. Soc. B, 366(1565), 688–696. doi:10.1098/rstb.2010.0200

Kriska, G., Bernáth, B., Farkas, R., & Horváth, G. (2009). Degrees of polarization of reflected light eliciting polarotaxis in dragonflies (Odonata), mayflies (Ephemeroptera) and tabanid flies (Tabanidae). J. Insect Physiol., 55(12), 1167–1173. doi:10.1016/j.jinsphys.2009.08.013

Kriska, G., Csabai, Z., Boda, P., Malik, P., & Horváth, G. (2006). Why do red and dark-coloured cars lure aquatic insects? The attraction of water insects to car paintwork explained by reflection-polarization signals. Proc. R. Soc. B, 273(1594), 1667–1671. doi:10.1098/rspb.2006.3500

Kriska, G., Horváth, G., & Andrikovics, S. (1998). Why Do Mayflies Lay their Eggs En Masse on Dry Asphault Roads? Water-Imitating Polarized Light Reflected from Asphault Attracts Ephemoptera. J. Exp. Biol., 201, 2273–2286.

Labhart, T. (1996). How polarization-sensitive interneurones of crickets perform at low degrees of polarization. J. Exp. Biol., 199(Pt 7), 1467–75.

Labhart, T. (2016). Can invertebrates see the e-vector of polarization as a separate modality of light?, 3844–3856. doi:10.1242/jeb.139899

Labhart, T., & Meyer, E. P. (1999). Detectors for polarized skylight in insects: a survey of ommatidial specializations in the dorsal rim area of the compound eye. Microsc. Res. Tech., 47(6), 368–79.

Lambrinos, D., Kobayashi, H., Pfeifer, R., Maris, M., Labhart, T., & Wehner, R. (1997). An Autonomous Agent Navigating with a Polarized Light Compass. Adapt. Behav., 6(1), 131–161. doi:10.1177/105971239700600104

Land, E. H. (1951). Some Aspects of the Development of Sheet Polarizers. J. Opt. Soc. Am., 41(12), 957–963. doi:10.1364/JOSA.41.000957

Lerner, A. (2014). Underwater Polarization by Scattering Hydrosols. In Polarized Light in Animal Vision (G. Horváth, Ed.). Berlin/Heidelberg: Springer. doi:10.1007/978-3-642-54718-8

Lind, O., Sunesson, T., Mitkus, M., & Kelber, A. (2012). Luminance-dependence of spatial vision in budgerigars (Melopsittacus undulatus) and Bourke’s parrots (Neopsephotus bourkii). J. Comp. Physiol. A, 198(1), 69–77. doi:10.1007/s00359-011-0689-7

Lythgoe, J. N., & Hemmings, C. C. (1967). Polarized Light and Underwater Vision. Nature, 213(5079), 893–894.

Marshall, N. J., Roberts, N. W., & Cronin, T. W. (2014). Polarisation signals. In Polarized Light in Animal Vision (G. Horváth, Ed.). Berlin/Heidelberg: Springer. doi:10.1007/978-3-642-54718-8

Maxwell, D. J., Partridge, J. C., Roberts, N. W., Boonham, N., & Foster, G. D. (2016). The Effects of Plant Virus Infection on Polarization Reflection from Leaves. PLoS ONE, 11(4), e0152836. doi:10.1371/journal.pone.0152836

Mäthger, L. M., Lohmann, K. J., Limpus, C. J., & Fritsches, K. A. (2011). An unsuccessful attempt to elicit orientation responses to linearly polarized light in hatchling loggerhead sea turtles (Caretta caretta). Phil. Trans. R. Soc. B, 366(1565), 757–762. doi:10.1098/rstb.2010.0212

McCann, G. D., & Arnett, D. W. (1972). Spectral and polarization sensitivity of the dipteran visual system. J. Gen. Physiol., 59(5), 534–58.

Melgar, J., Lind, O., & Muheim, R. (2015). No response to linear polarization cues in operant conditioning experiments with zebra finches. J. Exp. Biol., 218, 2049–2054. doi:10.1242/jeb.122309

Michelson A. A. (1911). On metallic colouring in birds and insects. Philos. Mag. 21, 554–567. doi:10.1080/14786440408637061

Mie, G. (1908). Beiträge zur Optik trüber Medien, speziell kolloidaler Metallösungen. Ann. Phys., 330(3), 377–445. doi:10.1002/andp.19083300302

Muheim, R. (2011). Behavioural and physiological mechanisms of polarized light sensitivity in birds. Phil. Trans. R. Soc. B, 366(1565), 763–771. doi:10.1098/rstb.2010.0196

Mussi, M., Haimberger, T. J., & Hawryshyn, C. W. (2005). Behavioural discrimination of polarized light in the damselfish Chromis viridis (family Pomacentridae). J. Exp. Biol., 208(16), 3037–3046. doi:10.1242/jeb.01750

Northmore, D. P. M., & Dvorak, C. A. (1979). Contrast sensitivity and acuity of the goldfish. Vision Res., 19, 255–261.

Park, H., & Crozier, K. B. (2015). Elliptical silicon nanowire photodetectors for polarization-resolved imaging. Opt. Express, 23(6), 7209–7216. doi:10.1364/OE.23.007209

Parkyn, D., Austin, J. D., & Hawryshyn, C. W. (2003). Acquisition of polarized-light orientation in salmonids under laboratory conditions. Anim. Behav., 65(5), 893–904. doi:10.1006/anbe.2003.2136

Pfeiffer, K., Negrello, M., & Homberg, U. (2011). Conditional perception under stimulus ambiguity: polarization-and azimuth-sensitive neurons in the locust brain are inhibited by low degrees of polarization. J. Neurophysiol., 105(1), 28–35. doi:10.1152/jn.00480.2010

Powell, S. B., & Gruev, V. (2013). Calibration methods for division-of-focal-plane polarimeters. Optics Express, 21(18), 579–592. doi:10.1364/0E.21.021039

Roberts, N. W., How, M. J., Porter, M. L., Temple, S. E., Caldwell, R. L., Powell, S. B., … Cronin, T. W. (2014). Animal Polarization Imaging and Implications for Optical Processing. Proc. IEEE, 102(10), 1427–1434. doi:10.1109/JPROC.2014.2341692

Rossel, S., Wehner, R., & Lindauer, M. (1978). E-Vector Orientation in Bees. J. Comp. Physiol. A, 125(1), 1–12. doi:10.1007/BF00656826

Sakura, M., Okada, R., & Aonuma, H. (2012). Evidence for instantaneous e-vector detection in the honeybee using an associative learning paradigm. Proc. R. Soc. B, 279(1728), 535–542. doi:10.1098/rspb.2011.0929

Schechner, Y. Y. (2011). Inversion by P4: polarization-picture post-processing. Phil. Trans. R. Soc. B, 366(1565), 638–648. doi:10.1098/rstb.2010.0205

Schwind, R. (1995). Spectral regions in which aquatic insects see reflected polarized light. J. Comp. Physiol. A, 177, 439–448.

Schwind, R. (1983). A polarization-sensitive response of the flying water bug Notonecta glauca to UV light. J. Comp. Physiol. A, 150(1), 87–91. doi:10.1007/BF00605291

Schwind, R. (1989). A Variety of Insects are Attracted to Water by Reflected Polarized Light. Naturwissenschaften, 76, 377–378.

Schwind, R. (1991). Polarization vision in water insects and insects living on a moist substrate. J. Comp. Physiol. A, 168, 531–540.

Schwind, R. (1999). Daphnia pulex swims towards the most strongly polarized light—a response that leads to “shore flight.”; J. Exp. Biol., 202, 3631–3635.

Sharkey, C. R., Partridge, J. C., & Roberts, N. W. (2015). Polarization sensitivity as a visual contrast enhancer in the Emperor dragonfly larva, Anax imperator. J. Exp. Biol., 218(21), 3399–3405. doi:10.1242/jeb.122507

Shashar, N., Hagan, R., Boal, J. G., & Hanlon, R. T. (2000). Cuttlefish use polarization sensitivity in predation on silvery fish. Vision Res., 40(1), 71–75.

Shashar, N., & Hanlon, R. T. (1997). Squids (Loligo pealei and Euprymna scolopes) Can Exhibit Polarized Light Patterns Produced by Their Skin. Biol. Bull., 193, 207–208.

Shashar, N., Rutledge, P., & Cronin, T. W. (1996). Polarization vision in cuttlefish in a concealed communication channel? J. Exp. Biol., 199, 2077–2084.

Shashar, N., Sabbah, S., & Aharoni, N. (2005). Migrating locusts can detect polarized reflections to avoid flying over the sea. Biol. Lett., 1(4), 472–475. doi:10.1098/rsbl.2005.0334

Shurcliff, W. A. (1955). Haidinger’s Brushes and Circularly Polarized Light. JOSA, 45(5), 399.

Stevens, M., Párraga, C. A., Cuthill, I. C., Partridge, J. C., & Troscianko, T. S. (2007). Using digital photography to study animal coloration. Biol. J. Linn. Soc., 90(2), 211–237. doi:10.1111/j.1095- 8312.2007.00725.x

Stewart, F. J., Kinoshita, M., & Arikawa, K. (2017). A Novel Display System Reveals Anisotropic Polarization Perception in the Motion Vision of the Butterfly Papilio xuthus. Integr. Comp. Biol., 1–9. doi:10.1093/icb/icx070

Strutt, J. W. (1871). On the Light from the Sky, its Polarization and Colour. The London, Edinburgh and Dublin Philosophical Magazine and Journal of Science, xxxvii, 107–120.

Taylor, D. H., & Adler, K. (1973). Spatial orientation by Salamanders using plane-polarized light. Science, 181(4096), 285–7.

Temple, S. E., Mcgregor, J. E., Miles, C., Graham, L., Miller, J., Buck, J., … Roberts, N. W. (2015). Perceiving polarization with the naked eye: characterization of human polarization sensitivity. Proc. R. Soc. B, 282(2015–0338). doi:10.1098/rspb.2015.0338

Temple, S. E., Pignatelli, V., Cook, T., How, M. J., Chiou, T.-H., Roberts, N. W., & Marshall, N. J. (2012). High-resolution polarisation vision in a cuttlefish. Curr. Biol., 22(4), R121–2. doi:10.1016/j.cub.2012.01.010

Tuthill, J. C., & Johnsen, S. (2006). Polarization sensitivity in the red swamp crayfish Procambarus clarkii enhances the detection of moving transparent objects. J. Exp. Biol., 209(9), 1612–6. doi:10.1242/jeb.02196

Wang, X., Gao, J., Fan, Z., & Roberts, N. W. (2016). An analytical model for the celestial distribution of polarized light, accounting for polarization singularities, wavelength and atmospheric turbidity. J. Opt., 18(6), 65601. doi:10.1088/2040-8978/18/6/065601

Wardill, T. J., Gonzalez-Bellido, P. T., Crook, R. J., & Hanlon, R. T. (2012). Neural control of tuneable skin iridescence in squid. Proc. R. Soc. B, 279(1745), 4243–4252. doi:10.1098/rspb.2012.1374

Wehner, R. (2001). Polarization vision—a uniform sensory capacity? J. Exp. Biol., 204, 2589–2596.

Wehner, R., & Strasser, S. (1985). The POL area of the honey bee’s eye: behavioural evidence. Physiol. Entomol., 10(3), 337–349.

Wolff, L. B. (1990). Polarization-Based Material Classification from Specular Reflection. IEEE Trans. Pattern. Anal. Mach. Intell., 12(11), 1059–1071.

Wolff, L. B. (1997). Polarization vision: a new sensory approach to image understanding. Image Vis. Comput., 15, 81–93. doi:10.1016/S0262-8856(96)01123-7

Wolf, R., Gebhardt, B., Gademann, R., & Heisenberg, M. (1980). Polarization Sensitivity of Course Control in Drosophila melanogaster. J. Comp. Physiol. A, 139(3), 177–191. doi:10.1007/BF00657080

York, T., Powell, S. B., Gao, S., Kahan, L., Charanya, T., Saha, D., … Gruev, V. (2014). Bioinspired Polarization Imaging Sensors: From Circuits and Optics to Signal Processing Algorithms and Biomedical Applications. Proc. IEEE, 102(10), 1450–1469. doi:10.1109/JPROC.2014.2342537

Yu, L., & Rawat, B. S. (1992). Mode-Coupling Analysis of Depolarization Effects in a Multimode Optical Fiber. J. Lightwave Technol., 10(5), 556–562. doi:10.1109/50.136088

